# β-Myosin inhibition in skeletal muscle: Mavacamten’ s impact on the diaphragm

**DOI:** 10.64898/2026.11.17.767324

**Authors:** Ayesha Sarfraz, Arnab Nayak, Tim Holler, Theresia Kraft, Mamta Amrute-Nayak

## Abstract

The diaphragm, a primary respiratory muscle, contains nearly equal proportions of slow, Type-1 and fast, Type-II muscle fibers. Muscle fiber types -classified based on their shortening velocities and force generation abilities -are largely determined by distinct myosin II isoforms. Mavacamten (MAVA), a small molecule inhibitor of β-cardiac myosin ATPase, is currently used for the treatment of obstructive Hypertrophic Cardiomyopathy (HOCM). Notably, β-myosin heavy chain (β-MyHC) is not restricted to the ventricular myocardium but is also expressed in slow fibers, raising concerns about off-target effects of therapy used to treat cardiomyopathic conditions. Given that nearly 50 % of the fibers express β-MyHC in the diaphragm, we hypothesize that Mava may affect diaphragm muscle activity. Using an in vitro motility assay, we investigated the effects of Mava on native myosins isolated from rabbit diaphragm muscle and measured the velocity of actin and regulated thin filaments. Mava caused a dose-dependent decrease in myosin-driven actin filament velocity, with an apparent IC_50_ of 1350 nM. Importantly, this inhibition was reversible upon drug washout, indicating a non-permanent modulatory effect. To determine the drug’s fiber-type specificity, we tested pure slow- and fast-myosin isoforms from the slow-twitch M. soleus and fast-twitch M. psoas. Both isoforms were inhibited by Mava, but slow myosin exhibited nearly 20-fold greater sensitivity than fast myosin, suggesting a potentially stronger impact on slow fibers. Interestingly, the slow myosin isoform showed even higher sensitivity to Mava than β-cardiac myosin. Further mechanistic analysis distinguished two modes of action: (i) promotion of OFF state of myosin, likely reducing a pool of active motors interacting with actin filaments, and (ii) modulation of the actomyosin biochemical cycle. Altogether, given the diaphragm’s reliance on both fiber types for sustained respiratory function, this off-target effect may impair respiratory performance in patients undergoing therapy, underscoring the need for respiratory function monitoring. The studies have direct implications for understanding the modulation of myosin function under physiological and pathophysiological conditions as well as following pharmacological interventions.

**Statement of significance:** By directly evaluating the effects of mavacamten (MAVA) on diaphragm myosin, this study bridges a critical mechanistic gap between cardiac myosin pharmacology and respiratory muscle physiology. Our findings show that MAVA strongly inhibits slow skeletal myosin activity, which accounts for nearly 50% of the diaphragm myosin population. In addition, we distinguished two distinct inhibitory modes of action for myosins in different conformational and functional configurations. This work establishes a framework for understanding MAVA’s potential off-target effects on respiratory mechanics and provides a foundation for future investigation into its consequences for diaphragm contractility in HCM patients undergoing MAVA treatment.

## Introduction

The diaphragm is the principal inspiratory skeletal muscle contracting rhythmically throughout life(*1*). It is innervated primarily by the phrenic nerve and is organized into motor units that differ in contractile speed, fatigue resistance, and force-generating capacity. As a mixed skeletal muscle, the diaphragm consists of fibers with diverse contractile and metabolic properties. Fiber-type classification is mainly based on myosin heavy chain (MyHC) isoform expression. Type I fibers express slow MyHC, are slow-twitch, oxidative, and highly fatigue-resistant. Type IIA fibers express MyHC2A, are intermediate fast-twitch fibers with substantial oxidative capacity, whereas Type IIX fibers express MyHC2X and have faster shortening kinetics and greater glycolytic capacity. Type IIB fibers express MyHC2B and are generally the fastest and most fatigable. This mixed fiber population enables the diaphragm to adapt to different physiological demands, functioning as an endurance muscle during quiet breathing and generating rapid, high forces during actions like coughing, sneezing, vomiting, and airway obstruction (*2*). The functional significance of fiber-type heterogeneity is particularly evident within motor units. Diaphragm motor units can be broadly classified as slow fatigue-resistant (S), fast fatigue-resistant (FR), fast fatigue-intermediate (FInt), and fast fatigable (FF), each associated with characteristic MyHC expression, twitch kinetics, force-generating properties, and resistance to fatigue (*2*). During normal breathing, only a fraction of the diaphragm’s total force-generating capacity is used, whereas airway occlusion, coughing, and sneezing require recruitment of additional, higher-force motor units (*1*). Reliance of quiet breathing mainly on fatigue-resistant motor units prevents unnecessary fatigue during lifelong breathing but allows rapid force escalation through more fatigable units when the respiratory system is challenged. Species differences are important when interpreting diaphragm fiber types: adult human skeletal muscle is generally dominated by type I, IIA, and IIX fibers, with little or no MyHC2B expression under normal conditions, whereas rodent diaphragms often contain the MyHC2B-positive fibers additionally (*3*, *4*). In adult humans, slow and fast fibers are present in approximately equal proportions in the diaphragm (*5*).

Myosin II is the principal molecular motor that determines the force-generating capacity of skeletal and cardiac muscle. As an ATP-dependent motor, it converts the chemical energy of ATP hydrolysis into mechanical force and movement through a cyclic process whereby myosin binds ATP, hydrolyses it to ADP and inorganic phosphate (Pi), interacts with actin, releases hydrolysis products Pi and ADP, associated with force-producing conformational changes (the generation of powerstroke), and then detaches from actin to reset the cycle (*6*, *7*). Myosin II is a hexameric complex composed of two myosin heavy chains, two essential light chains (ELCs), and two regulatory light chains (RLCs). Each heavy chain contains an N-terminal globular motor domain with ATP- and actin-binding sites, a neck or lever-arm region that binds the light chains, and a long α-helical coiled-coil tail that mediates dimerization and contributes to thick-filament assembly. The myosin heavy chain determines motor properties including actin-activated ATPase activity, duty ratio, ADP release, actin affinity, force production, and shortening velocity(*6*, *8*). The ELCs and RLCs stabilize the lever arm and modulate myosin function by fine-tuning actomyosin attachment/detachment kinetics, force production, and mechanical efficiency. Together, these structural and functional components enable the myosin motor to drive muscle contraction.

In the human and higher mammals heart, the β-cardiac myosin heavy chain (β-MyHC), encoded by the *MYH7* gene, is the dominant ventricular isoform (*9*). It exhibits slower ATPase activity and actin-sliding speed compared to the α-cardiac myosin isoform, which is predominantly expressed in the atrium. Notably, β-MyHC is also expressed in slow-twitch type I skeletal muscle fibres (*3*). β-cardiac myosin can adopt an autoinhibited ‘OFF-state’ conformation, also known as the interacting-heads motif (IHM) state, whereby the two myosin heads are folded back and sequestered onto the proximal S2 tail region of the thick filament (*10*–*14*). This structural arrangement reduces ATP turnover and limits myosin-head availability for actin binding, contributing to a low energy consuming state. Although the biochemical super relaxed state (SRX) - characterized by low ATP turnover is widely studied, its direct correspondence to the IHM state remains under investigation. The modulation of β-cardiac myosin function in a sarcomere therefore depends on the catalytic properties of individual myosin heads and the dynamic equilibrium between sequestered (OFF) and actin-accessible (ON) states. This balance is central to some of the current models of hypertrophic cardiomyopathy (HCM) pathogenesis. One of the disease model suggests that HCM -causing mutations in *MYH7* may increase myocardial contractile ability and force by altering the ATPase activity of individual heads or by releasing a larger fraction of heads from the sequestered OFF state (*15*). HCM, one of the most common inherited cardiomyopathies, is characterized by left ventricular hypertrophy, often with asymmetric septal thickening (*16*–*18*). It is considered a ‘disease of the sarcomere’ and a leading cause of sudden cardiac death (SCD) in young athletes. In obstructive HCM (HOCM), left ventricular outflow tract obstruction arises from a combination of septal hypertrophy, altered ventricular geometry, hyperdynamic systolic contraction, and systolic anterior motion of the mitral valve.

Currently, Mavacamten (MYK-461), a first-in-class, small-molecule inhibitor of cardiac myosin ATPase is approved for symptomatic obstructive HCM. Unlike conventional negative inotropes, it is a myotrope, acting directly by allosterically modulating its function to reduce contractility (*19*). The therapeutic rationale for MAVA is based on the observations that altered or increased sarcomeric force generation, increased velocity, high ATP turnover rate and increased Ca^2+^ sensitivity or so called hypercontractility contributes to HCM pathophysiology (*20*–*23*). Note that some studies found reduced sarcomeric force or hypocontractility in selected HCM mutation-bearing sarcomeric proteins (*24*–*26*). Preclinical studies demonstrated that MAVA reduces cardiac myosin ATPase activity and attenuates hypertrophic remodeling in HCM mouse models (*19*). Mechanistically, it stabilizes myosin in low-ATPase, super-relaxed (SRX) states, reducing the population of heads available for productive actin interaction by affecting the transition from non-force-generating to force-generating actomyosin states(*27*, *28*). The EXPLORER-HCM trial demonstrated MAVA’s clinical efficacy, showing significant improvement in exercise capacity, post-exercise left ventricular outflow tract (LVOT) gradient, and New York Heart Association (NYHA) functional class in patients with symptomatic obstructive HCM) (*29*). The VALOR-HCM trial further demonstrated that Mavacamten reduced the proportion of patients eligible for septal reduction therapy-such as alcohol septal ablation or septal myectomy among those with severe symptomatic disease. Long-term data from VALOR-HCM confirm sustained improvement in obstruction and symptoms during continued treatment. However, careful monitoring remains essential as excessive myosin inhibition may impair systolic function (*30*). MAVA exhibits preferential inhibition of cardiac myosin with significant effects on β-cardiac isoform observed at clinically relevant concentrations, while fast myosin is inhibited only at higher concentrations (*31*). This differential potency supports the compound’s cardiac selectivity, although it does not completely exclude off-target effects on skeletal muscle (*31*–*33*). This distinction is particularly important in the context of systemic pharmacology, as oral administration exposes MAVA to tissues beyond the myocardium, necessitating careful evaluations of potential effects, especially during long-term therapy.

The potential link between diaphragm function and MAVA arises from a fundamental physiological fact, i.e., slow-twitch type-I fibers expressing β-MyHC constitute a substantial proportion of the diaphragm’s contractile function. While MAVA has been extensively characterized in cardiac myosin, HCM models, and patients, its direct effects on diaphragm myosin remain unexplored. Given that β-MyHC is expressed in both cardiac and slow-twitch Type-1 fibers, and that MAVA selectively inhibits β-MyHC at clinically relevant concentrations, we hypothesize that systemic exposure to MAVA may also impair diaphragm myosin function and thereby respiratory function. Here, we systematically evaluated MAVA’s effects on the diaphragm, M. psoas, M. soleus and cardiac myosin function and recovery after removal of the drug from the measurement chamber. We found that slow myosin isoform exhibits even greater sensitivity to MAVA inhibition than β-cardiac myosin. Remarkably, we resolved MAVA’s dual mode of inhibitory action, (1) on myosin heads adopting distinct conformations in the actin-unbound state and (2) impeding kinetic transition within actomyosin cross-bridge cycle, resulting in overall reduction in actin gliding speed. The varying degrees of functional inhibition and recovery observed across myosin isoforms suggest that MAVA’s effect on the muscle’s contractile function will be determined by their fiber type composition, and especially the muscles such as diaphragm’s performance will be more susceptible to MAVA.

## Results

### Diaphragm composition

We first evaluated the diaphragm muscle composition for its fiber type and myosin isoform content. The diaphragm is anatomically subdivided into sternal, costal, and crural regions based on the origin of muscle fibers(*2*). Here, we used the muscle tissue isolated from costal region of the rabbit diaphragm (Fig. 1A), as it represents the primary contractile portion of the respiratory muscle and provides a consistent and representative sampling site (See Fig. S1 for original rabbit diaphragm muscle).

**Fig. 1.**
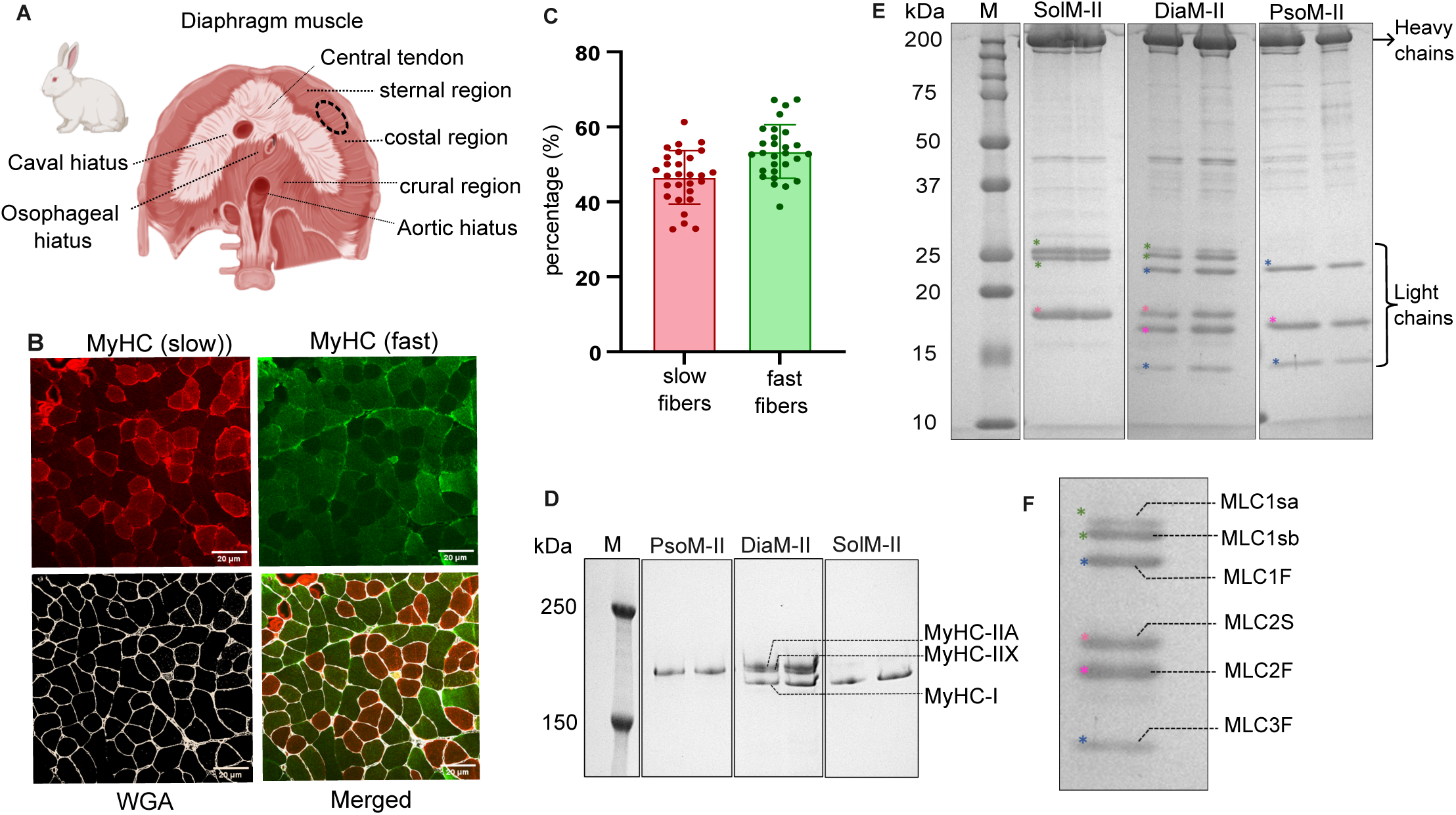
Characterization of rabbit diaphragm. **(A)** Schematic representation of the rabbit diaphragm showing the sternal, costal, and crural regions. The encircled region is the site of tissue collection for our experiments. **(B)** Representative transverse cryosections immunolabelled for slow myosin heavy chain (slow MyHC; red) and fast myosin heavy chain (fast MyHC; green). Wheat germ agglutinin (WGA; white) delineates individual muscle-fibre boundaries; the merged image shows the mosaic distribution of slow- and fast-MyHC-positive fibres. **(C)** Quantification of the percentages of slow- and fast-twitch fibres in the analysed diaphragm sections. Bars represent mean ± SD, with individual measurements overlaid. Number of cryosections = 28, Number of rabbits = 2, each spherical symbol represents one cryosection. P = 0.0007, unpaired two-tailed Student’s t-test. **(D)** Representative gel-electrophoretic separation of myosin heavy-chain isoforms in PsoM-II, DiaM-II and SolM-II. PsoM-II was analysed at 0.75 and 1.5 µg, DiaM-II at 2.5 and 5.0 µg, and SolM-II at 0.75 and 1.5 µg. Only the lower part of the gel is shown to highlight the myosin heavy chains. DiaM-II contains one slow MyHC-I band together with two fast MyHC bands corresponding to MyHC-IIA and MyHC-IIX. M, molecular-mass marker.**(E)** Representative SDS-PAGE analysis of diaphragm myosin II (DiaM-II), soleus myosin II (SolM-II) and psoas myosin II (PsoM-II). The upper and lower brackets indicate the myosin heavy-chain and light-chain regions, respectively. SolM-II contains the characteristic slow skeletal light-chain isoforms MLC1sa, MLC1sb and MLC2S, whereas PsoM-II contains the fast skeletal isoforms MLC1F, MLC2F and MLC3F. **(F)** DiaM-II contains bands corresponding to both slow and fast myosin light-chain isoforms, consistent with its mixed fibre-type composition. Coloured asterisks indicate the assigned light-chain bands.

To assess fiber-type composition, cryosections of the diaphragm tissue were generated. Both transverse and longitudinal sections from the costal region were subjected to immunofluorescence staining for myosin heavy chain (MyHC) isoforms. Slow and fast fibers expressing MyHC-slow and MyHC-fast, respectively, were probed using isoform-specific antibodies (Fig.1B). Wheat germ agglutinin (WGA) staining was used to mark muscle fiber boundaries by binding to membrane-associated glycoproteins (*34*). A clear mosaic distribution of slow- and fast-twitch fibers was evident in the cryosections. Notably, no co-expression of fast and slow MyHCs was evident in single fibers, as reported for several species (*35*). Fiber type analysis based on the myosin heavy chain isoforms confirmed that the rabbit diaphragm contains both slow- and fast-twitch fibers with nearly equal proportion (Fig. 1C).

To characterize the myosin composition in diaphragm, myosin heavy chain (MyHC) and light chain (LC) isoforms were analyzed by SDS-PAGE. Rabbit *M. soleus* which is predominantly composed of slow fibers (up to 97%) (*36*), and rabbit *M. psoas* enriched in fast-twitch fibers (*36*), were used as references for slow and fast myosin isoforms. As expected, SolM-II predominantly expressed slow MyHC isoform, while PsoM-II primarily expressed fast MyHC isoform. In comparison, the DiaM-II contained a slow (MyHC-I) and two fast MyHC isoforms (MyHC-IIA, MyHC-IIX) as shown in Fig. 1D. Fig. 1E and F shows the analysis of myosin light chain (MLC) isoforms. Distinct MLC isoforms corresponding to slow (MLC1sa, MLC1sb, MLC2s) and fast (MLC1F, MLC2F, MLC3F) fiber types were clearly resolved in the control samples (Fig. 1E–F). The diaphragm myosin exhibited a combination of all identified MLC isoforms, consistent with contributions from both slow- and fast-twitch fibers (Fig. 1F)

### Mavacamten affects DiaM-II function

Using standard actin filament gliding assay (*37*) as schematized in Fig. 2A, we evaluated the actin filament gliding speed driven by myosin isolated from diaphragm tissue (Fig.2B). As control, we first assessed whether DMSO used as solvent for Mava has influence on myosin activity. We chose the highest concentration of DMSO that is used in our experiments, i.e., 1% of volume to test its effect. As shown in Fig. 2B, no significant difference in actin sliding velocity was observed between the control condition at 2 mM ATP and the DMSO-treated samples (p = 0.072), indicating that DMSO does not affect DiaM-II activity under these conditions (Fig. 2B). We assessed two main aspects of MAVA’s effects on myosin function;1) the functional change after exposure to MAVA under ‘treatment’ and the ‘recovery’ after removal of MAVA and motility measurements under normal condition of 2 mM ATP as schematized in Fig. 2C. The effect of MAVA on DiaM-II function was then evaluated over a wide range of MAVA concentrations (1 nM to 100 µM). A dose-dependent decrease in actin filament sliding velocity was observed with increasing MAVA concentration (Fig. 2D). The dose–response curve yielded an IC_50_ of about 1.35 µM. Native thin filaments (nTFs) are physiological functional partners of myosins, therefore in the next step, we investigated effects of MAVA on nTFs motility. nTFs were isolated from diaphragm muscle (Fig. S2) and assessed for their composition of regulatory components tropomyosin and troponins (Tm-Tn), which constituted a mixture of slow and fast fiber characteristics. Ca^2+^-mediated activation of the regulation nTFs was verified by comparing the motility under activating conditions (pCa 4.0) against the low Ca^2+^ conditions of pCa 8.0 (Fig. S3A). nTF motility driven by DiaM-II in the presence of Mava was evaluated. Interestingly, DiaM-II interacting with nTFs exhibited greater sensitivity to MAVA, as indicated by a lower IC₅₀ value compared to bare actin filaments, i.e., 691 nM and 1.35 µM, respectively (Fig. S3B-C). These findings suggest that mavacamten may influence nTF thin filament activation through its effect on diaphragm myosin function.

**Fig. 2:**
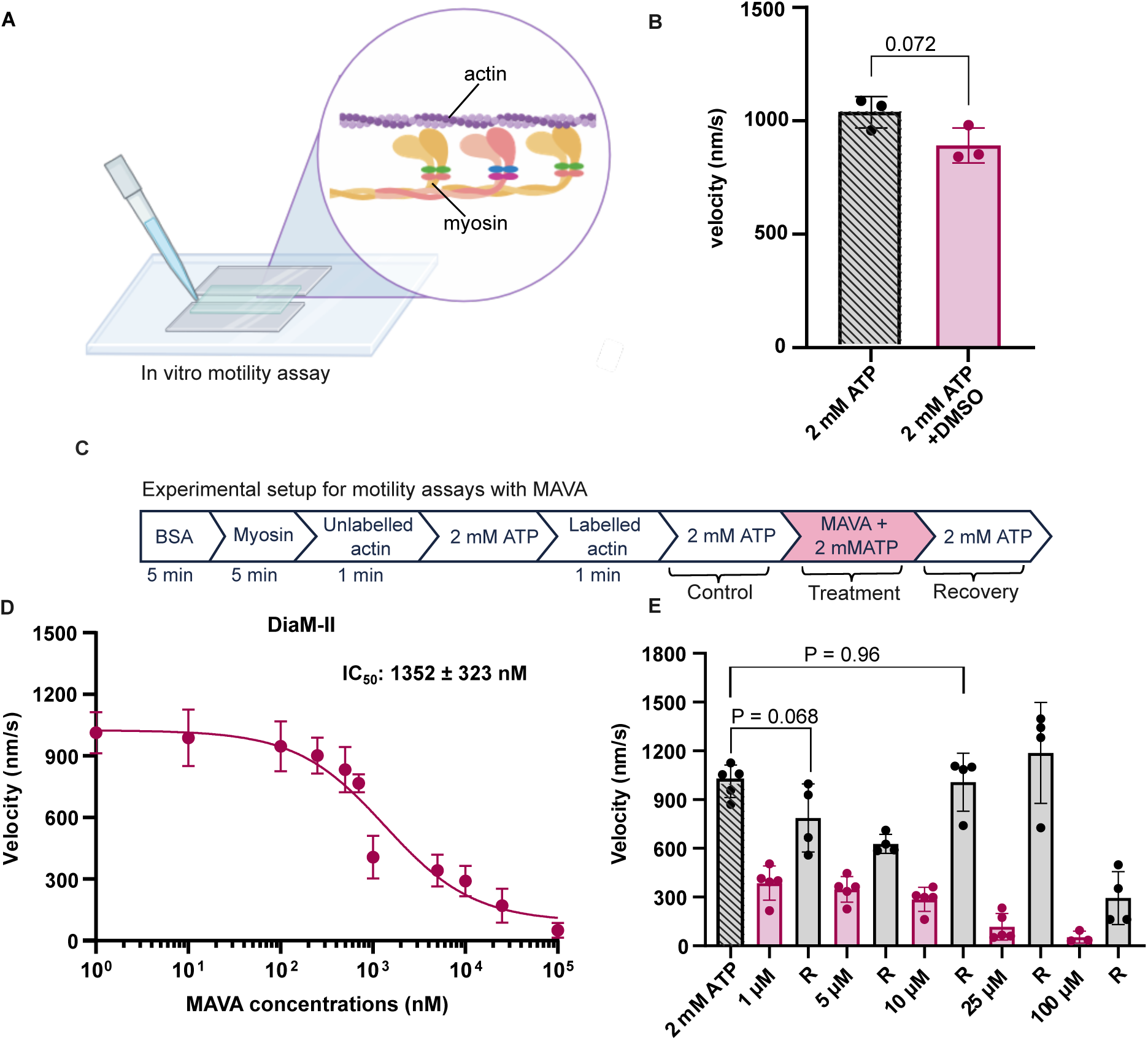
Functional analysis of DiaM-II activity and its modulation by MAVA. (A) A schematic representation of an in vitro motility assay. **(B)** Control experiment assessing the effect of DMSO on myosin activity. The highest solvent concentration used in the experiments, 1% (v/v) DMSO. Each data point represents a chamber, N=3. The number of actin filaments analyzed per condition; n = 100-120. **(C)** The experimental setup; flow cell recorded for control condition, MAVA ‘treatment’ and ‘recovery’ after MAVA washout. **(D)** MAVA concentration-dependent inhibition of rabbit diaphragm myosin**-**driven F-actin motility over the range of 1 nM–100 µM. The concentration–response was fitted by nonlinear regression using a four-parameter logistic model with a variable slope, yielding an IC_50_ of 1352 ± 323 nM. Each data point is an average of 3-5 individual chambers. **(E)** Recovery of DiaM-II activity following MAVA washout. DiaM-II-driven actin-filament velocity measured under the 2 mM ATP control condition, after treatment with 1, 5, 10, 25 or 100 µM MAVA with 2 mM ATP, and after removal of the drug by washing with assay buffer containing 2 mM ATP. R denotes the corresponding recovery measurement. Experiments were performed at 2 mM ATP and 22°C. N = 2-5, n = 100-350. Error bar: mean ± SD. Statistical significance was assessed using an unpaired two-tailed Student’s t-test.

### Myosin function recovery after Mava treatment

One of the critical questions is how well the myosin recovers post Mava treatment. To mimic this, we washed MAVA from the flow cells and studied the recovery. Following the recording at specified Mava concentration, the drug was washed out by thoroughly washing the motility chamber with 3x chamber volumes of assay buffer containing 2 mM ATP. ‘Recovery’ of actin sliding velocity was observed across the tested concentrations (1 µM, 5 µM, 10 µM, 25 µM, and 100 µM), confirming that MAVA-mediated inhibition is reversible in *in vitro* motility system (Fig. 2E). Interestingly, we also observed that higher the concentration of MAVA introduced in the motility chamber, better was the recovery after MAVA washout as observed at 10 µM and 25 µM. At very high concentrations of Mava, e.g. 100 µM this recovery trend was no longer observed. Note that the recovery was recorded 1 minute after Mava washout.

To determine whether incomplete recovery of actin filament gliding speed after Mava washout was due to insufficient equilibration time, we performed time-dependent recovery experiments at 1 µM, 10 µM, and 100 µM MAVA (Fig. S4). Extending the post washout incubation period up to 20 minutes did not yield significant improvement in actin filament sliding velocity across all tested MAVA concentrations. This lack of time-dependent recovery indicates that the residual inhibition is not attributable to incomplete drug removal from the assay chamber. Rather, a fraction of myosins may not reach full activation or cannot regain the pre-mava conformation to support normal myosin function at the tested Mava concentrations.

### Differential effects of MAVA on slow and fast myosin isoforms

As the diaphragm muscle has a mixture of slow- and fast-myosin isoforms (Fig. 1D-F), the observed inhibitory effect of MAVA on actin sliding speed raised the question whether both myosin isoforms are equally affected or one is preferentially inhibited. To address this, we considered three mechanistic scenarios that might explain our observations, as schematized in Fig. 3A, (i) selective inhibition of slow myosin, which would leave fast myosin activity intact and potentially enhance actin sliding velocity due to unhindered fast-myosin driven motility; (ii) inhibition of slow myosin, leading to either prolonged actomyosin dissociation or weak actomyosin interaction, which could generate mechanical drag on fast myosin driven movement, and thereby reduce overall gliding speed; or (iii) inhibition of both slow and fast myosin isoforms, resulting in reduction of the actin sliding speed.

**Fig. 3.**
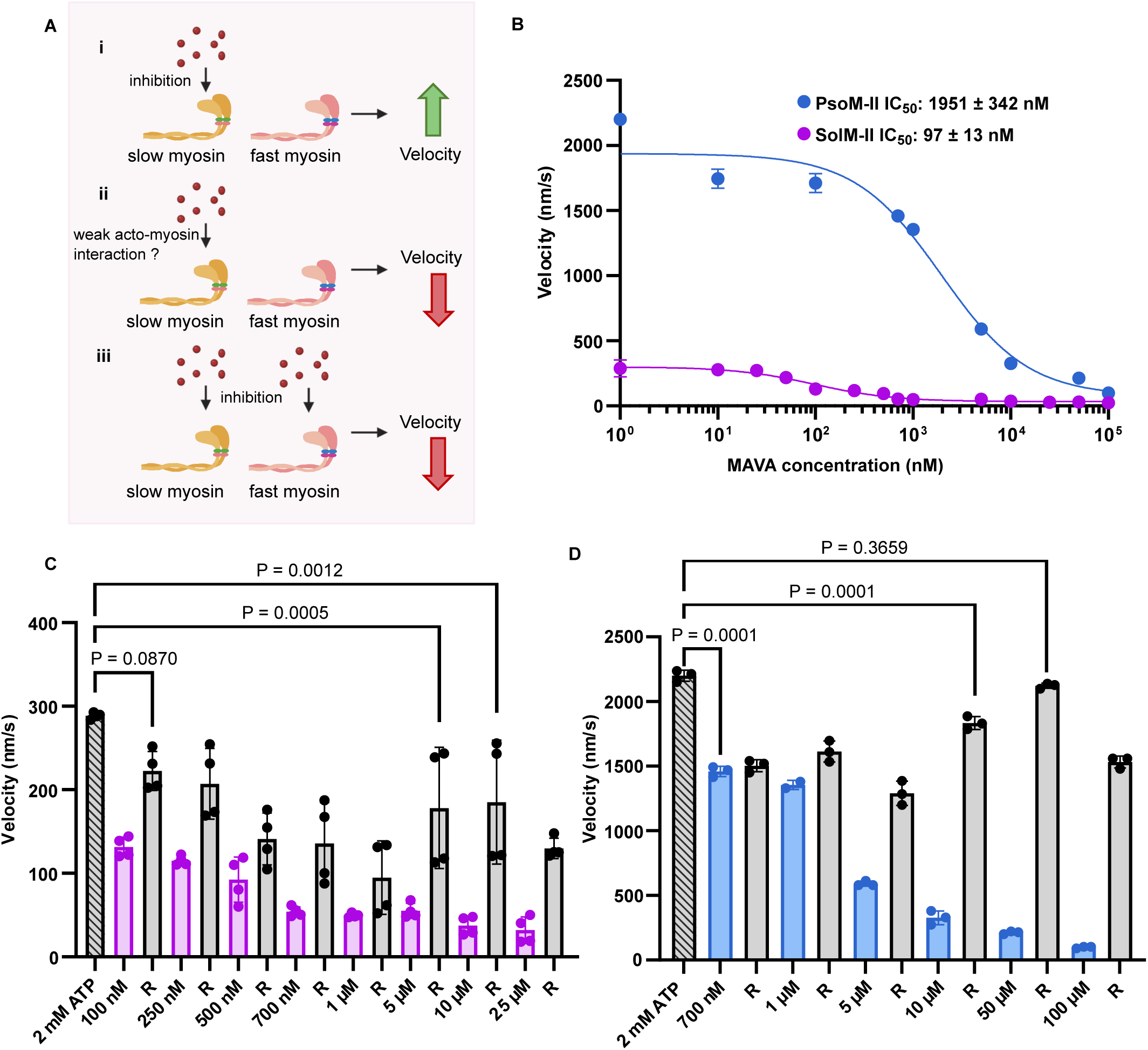
MAVA differentially modulates slow and fast skeletal myosin isoforms. **(A)** Schematic representation of three models considered for a mixed slow and fast myosin system, (i) selective inhibition of slow myosin with an increase in fast-myosin-driven velocity; (ii) inhibition of slow myosin accompanied by prolonged actomyosin attachment and drag on fast-myosin-driven movement or (iii) direct inhibition of both slow and fast myosin populations. **(B)** Comparison of MAVA concentration–response curves for SolM-II- and PsoM-II-driven actin-filament motility. Nonlinear regression fitting using a four-parameter logistic model with variable slope yielded IC_50_ values of 97 ± 13 nM for SolM-II and 1,951 ± 342 nM for PsoM-II. Note that for some of the velocity values, the error bars are smaller than the symbol values and therefore not visible. Mean velocities for each condition were estimated by fitting the Gaussian distribution. **(C)** Recovery of slow skeletal myosin following MAVA treatment. SolM-II velocity measured under the 2 mM ATP control condition and indicated MAVA concentrations with treatment (pink bars) and recovery R (grey bars) after exposure to specific MAVA concentration. **(D)** Recovery of fast skeletal myosin following MAVA treatment. PsoM-II velocity measured after MAVA treatment (Blue bars) and recovery R (grey bars). Each MAVA condition (treatment and recovery) was tested in a separate chamber. Experiments were performed at 2 mM ATP and 22°C. N = 2-4, n = 100-250. Error bar: mean ± SD. Statistical significance was assessed using an unpaired two-tailed Student’s t-test for selected comparisons.

The first assumption was ruled out as MAVA treatment led to a decrease –not an increase-in actin sliding velocity. To directly evaluate the second and third possibilities, the effects of MAVA were assessed on purified slow (SolM-II) and fast (PsoM-II) myosin isoforms isolated from *M. soleus* (predominantly slow-twitch fibers) and *M. psoas* (predominantly fast-twitch fibers), respectively. Under identical experimental conditions, Mava induced a clear dose-dependent reduction in actin filament sliding velocity for both SolM-II and PsoM-II across a range of Mava concentrations (Fig. 3B). Notably, the two isoforms exhibited distinct sensitivities to MAVA: the IC_50_ for SolM-II myosin was approximately 100 nM, whereas PsoM-II myosin showed a significantly higher IC_50_ of ∼2 µM, indicating that slow myosin is about 20-fold more sensitive to the drug than fast myosin isoforms. This differential sensitivity provides a mechanistic explanation for the observed inhibition in DiaM-II. The overall inhibitory effect of MAVA on DiaM-II motility likely arose from combined inhibition of both slow and fast myosin populations, rather than selective targeting of a single isoform.

To further investigate the nature of inhibition, the recovery of myosin activity following drug washout was assessed. In both SolM-II (Fig. 3C) and PsoM-II (Fig. 3D), actin sliding velocity was restored after removal of MAVA across all tested concentrations, suggesting that the inhibitory effect is reversible. A similar trend to that observed in DiaM-II (Fig. 2E) was evident, higher MAVA concentrations such as 10 µM and 50 µM led to more pronounced recovery of PsoM-II activity. Notably, in SolM-II, a partial recovery was observed at all the tested MAVA concentrations and the full restoration was not observed at any of the tested drug concentration, as observed for DiaM-II and PsoM-II. Collectively, these findings indicate that MAVA exerts a dose-dependent and reversible inhibitory effect on both slow and fast myosin isoforms, with SolM-II exhibiting a greater sensitivity to Mava.

### Enhanced sensitivity of SolM-II to MAVA compared to βM-II myosin

Mava is currently approved for the treatment of hypertrophic cardiomyopathy due to its selective modulation of beta cardiac myosin (βM-II) function. To elucidate the molecular basis of MAVA’s specificity, we compared the sensitivity of βM-II to that of SolM-II, which shares the same myosin heavy chain (MyHC) encoded by the *MYH7* gene and are endowed with the same light chain isoforms (MLC1sb/v and MLC2). However, slow-twitch skeletal muscle fibres uniquely expresses an additional essential light chain isoform, MLC1sa, which is absent in myocardium (*38*, *39*) (Fig. 4A). Given this difference, we investigated whether MLC1sa influences the sensitivity of SolM-II to MAVA. Dose–response analysis revealed a significant difference in the drug sensitivity of two myosin types (Fig. 4B). The IC_50_ for SolM-II myosin was ∼2-fold lower than that of βM-II myosin (IC_50_ of 97 nM *vs* 186 nM), indicating that SolM-II is more sensitive to MAVA inhibition. The only distinguishing feature between SolM-II and βM-II is the MLC1sa in SolM-II, strongly suggesting that the essential light chain isoform –MLC1sa-plays a modulatory role in determining myosins responsiveness to MAVA.

**Fig. 4.**
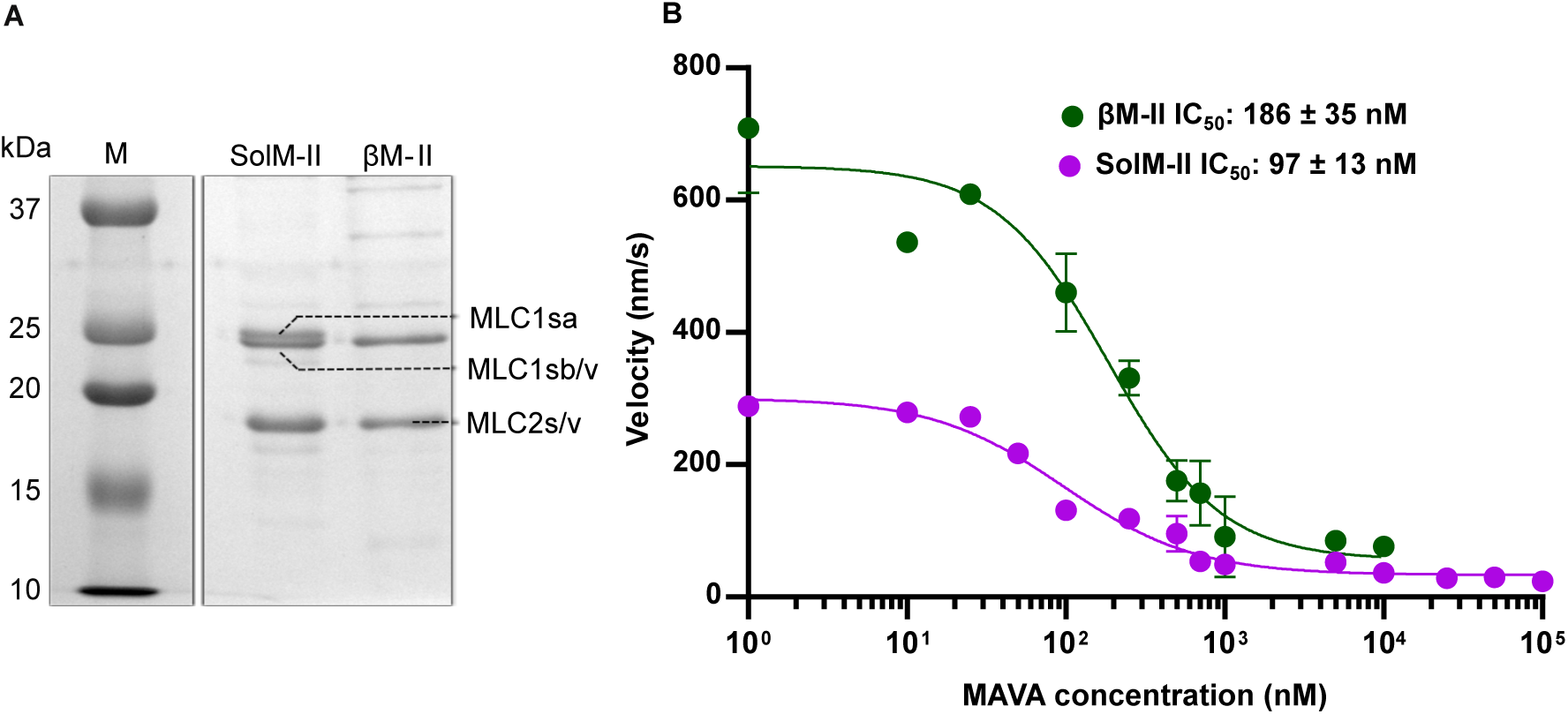
MAVA sensitivity of slow skeletal- and β-cardiac myosin. **(A)** Representative SDS-PAGE analysis of SolM-II and β-cardiac myosin II (βM-II). Both preparations contain MLC1sb/v and MLC2s/v, whereas the additional slow skeletal essential light-chain isoform MLC1sa is present in SolM-II but not in βM-II. M, molecular-mass marker. **(B)** Concentration–response curves for MAVA inhibition of SolM-II- and βM-II-driven actin-filament motility. Nonlinear regression fitting using a four-parameter logistic model with variable slope were used to obtain IC_50_ values of 97 ± 13 nM for SolM-II and 186 ± 35 nM for βM-II. Experiments were performed at 2 mM ATP and 22°C. Data is presented as mean ± SD; solid lines represent nonlinear dose–response fits. N = 3 (per condition), n = 150-200. Error bar: mean ± SD. Mean velocities for each condition were estimated by fitting the Gaussian distribution.

A comparative analysis of dose-response curves across different myosin isoforms (Fig. 5A) for their sensitivities towards Mava revealed following order: SolM-II > βM-II > DiaM-II > PsoM-II. SolM-II exhibited the highest sensitivity with an IC_50_ of about 97 nM, while the PsoM-II showed a lowest sensitivity with an IC_50_ ∼2 µM. DiaM-II displayed an intermediate IC_50_ ∼1.4 µM, consistent with its mixed composition of slow- and fast-M-II (Table 1). Assessment of myosin function recovery following MAVA-mediated inhibition (Fig. 5B) revealed that the extent of functional recovery varied significantly across all myosin isoforms. Near complete recovery was observed at intermediate Mava concentrations, consistent with transient and reversible Mava binding for DiaM-II, βM-II and PsoM-II. However, at very high concentrations of Mava such as ∼ 100 µM for PsoM-II and 25 µM for DiaM-II, myosin activity could not be recovered (Fig. 5B), suggesting residual inhibition persisted at high doses even after drug removal. Notably, SolM-II exhibited only partial recovery at all tested Mava concentrations.

**Fig. 5.**
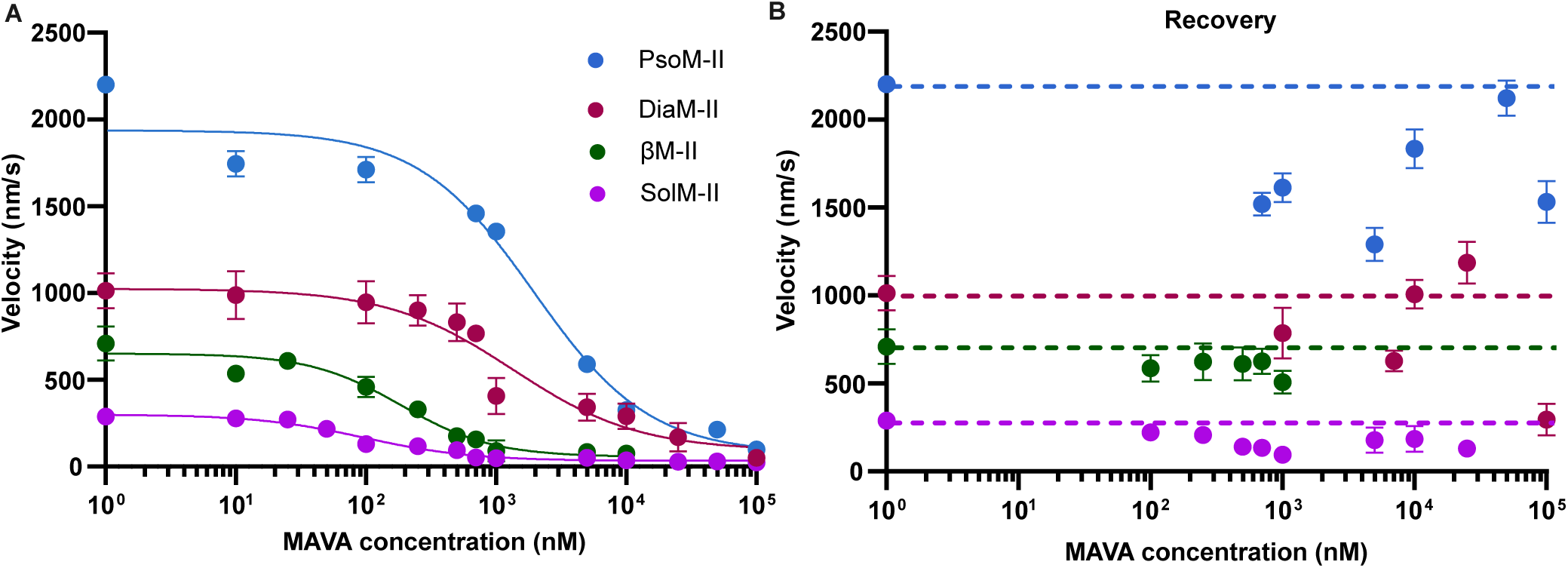
Comparison of myosin activities following MAVA treatment and washout. (A) Comparison of MAVA concentration–response curves for SolM-II, βM-II, DiaM-II and PsoM-II. (B) Recovery of actin filament velocities for different myosin isoforms, after exposure to indicated MAVA concentration. Dashed horizontal lines indicate the velocity at 2 mM ATP (control) for each myosin type. Each symbol represents the mean velocities ± SD. N = 2-5, n = 100-250. Mean velocities for each condition were estimated by fitting a Gaussian distribution.

**Table 1.** Summary of Vmax and IC₅₀ values for MAVA across rabbit myosin isoforms.

| **Myosin** | **Vmax<br>(nm/sec) ± SD** | **MAVA<br>IC<sub>50</sub> (nM)** |
|---|---|---|
| DiaM-II | 1013 ± 100 | 1352 ± 323 |
| PsoM-II | 2200 ± 43 | 1951 ± 342 |
| SolM-II | 288 ± 4 | 96.96 ± 13 |
| βM-II | 709 ± 98 | 186.1 ± 35 |

Mechanistically, MAVA has been shown to stabilize the interacting-heads motif (IHM) state of myosin-a structural conformation associated with myosin inhibition (*32*) and to promote a biochemical super-relaxed (SRX) state, characterized by slow ATP turnover (*40*). While the structural IHM and the SRX states are frequently used interchangeably in the literature, there is currently no consensus on their equivalence (*41*–*43*). Additionally, MAVA has been shown to reduce the rate of phosphate (Pi) release during the actomyosin cross-bridge cycle (*31*). Therefore, in the next step, we aimed to distinguish these two inhibitory modes of action via Mava.

### Mechanism of action of MAVA

To elucidate the molecular mechanisms underlying Mava’s observed inhibitory effects, we sought to distinguish between its impact on myosins structural or conformational state versus its influence on the kinetics of actomyosin crossbridge cycle. To isolate the possible effects on different states, we generated myosin subfragment-1 (S1), from PsoM-II using papain digestion (Fig. 6A and S5A). The resulting PsoS1 comprising a single-headed motor domain and lever arm, precludes the formation of the sequestered, IHM state. Therefore, any inhibition observed in PsoS1 is expected to arise from direct modulation of the actomyosin biochemical cycle rather than structural ‘OFF’ state, and thus providing an insight into MAVA’s action.

**Fig. 6.**
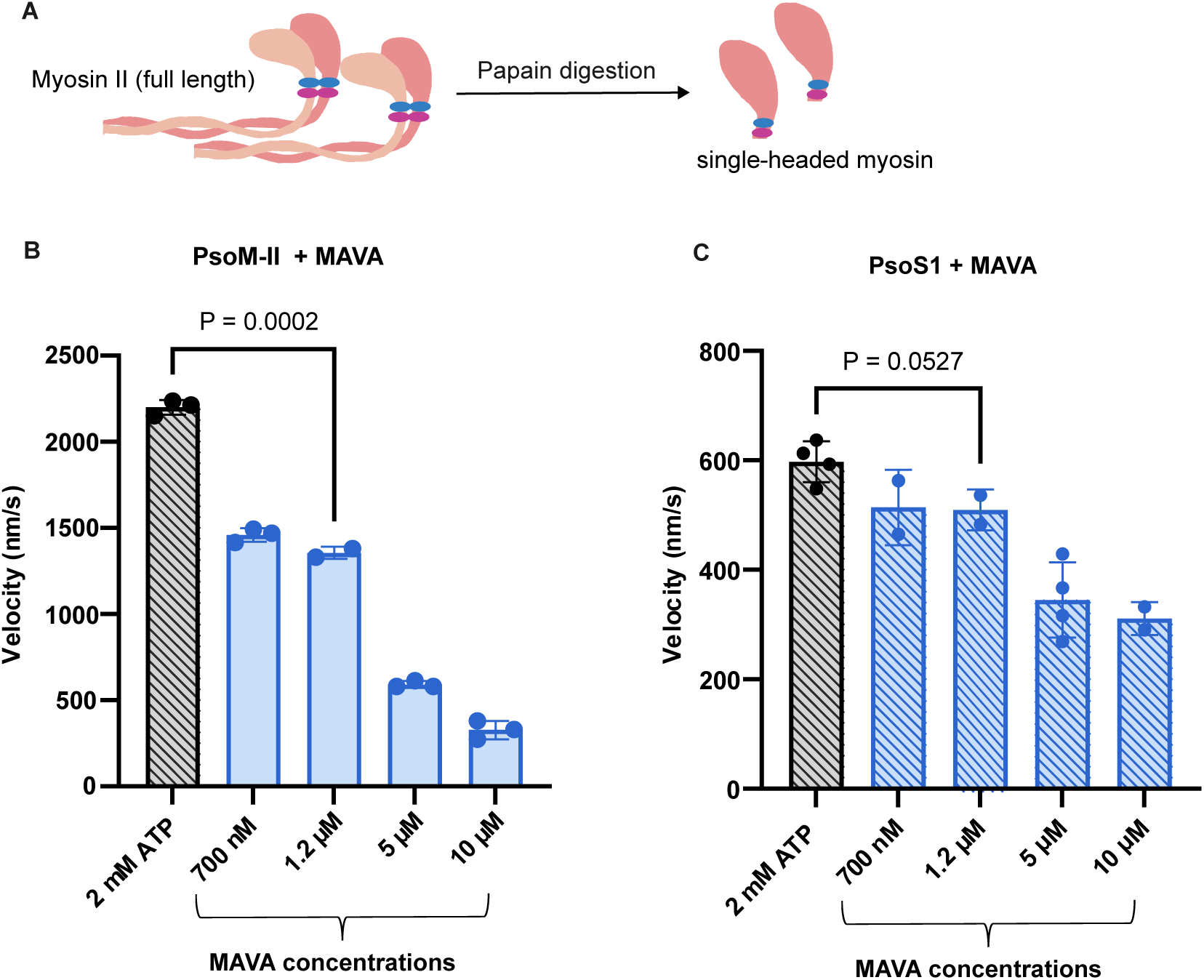
Differential effects of MAVA on full-length PsoM-II and single-headed PsoS1. **(A)** Schematic representation of the generation of myosin subfragment-1 (PsoS1) from full-length rabbit psoas myosin II (PsoM-II) by papain digestion. PsoS1 contains a single myosin motor domain and a lever arm with associated light chains (blue: ELC, pink: RLC). **(B, C)** Effects of increasing MAVA concentrations on actin-filament sliding velocity driven by full-length PsoM-II **(B)** and PsoS1 **(C)**. Statistical significance was assessed using an unpaired two-tailed Student’s t-test for selected comparisons. Bars represent mean ± SD. N= 2-4, n = 100 -150. Experiments were performed at 2 mM ATP and 22 °C.

As shown in (Fig. 6B and C), Mava inhibited the PsoM-II –driven actin gliding speed to a greater extent than PsoS1. For example, at a concentrations of 1.2 µM MAVA, PsoM-II mediated motility was reduced by ∼ 37 % of the control condition without drug (i.e., from 2200 nm/s to 1400 nm/s). In contrast, PsoS1-driven motility was only modestly affected, declining by about ∼17% (i.e., from 600 nm/s to 500 nm/s). Although a dose-dependent reduction in the activity was still present in PsoS1, the magnitude of inhibition was consistently and substantially lower across all Mava concentrations compared to full-length PsoM-II, indicating MAVA’s more pronounced impact on full-length myosin.

### MAVA induced inhibition of myosin

To further dissect the mechanistic basis of MAVA’s action, we investigated its effect on myosin in two distinct states: before binding to actin (i.e., in the absence of actin) and after engaging with actin. Note that, in the earlier experimental setups, Mava was added after myosin was already bound to actin filament in the last step, which may limit the ability to assess its impact on early modulatory steps. Importantly, the actin-unbound state of myosin is physiologically relevant, as the actomyosin interaction is strictly regulated by Ca^2+^ -dependent thin filament activation. In other words, in relaxed muscle, under low Ca^2+^ condition, Mava can access the myosin before its interaction with actin. This approach allowed us to probe MAVA’s myosin state-dependent mode of action.

We implemented a modified experimental design for the motility assay, as illustrated in Fig. 7A. The **Setup 1,** represents a standard motility assay, where myosin was first immobilized on a chamber surface, followed by the addition of F-actin (first unlabelled and then labelled), and motility was recorded at 2 mM ATP. After establishing baseline actin gliding velocity, MAVA was added to the chamber to assess its inhibitory effect, followed by drug washout to evaluate myosin function recovery. In **Setup 2**, we introduced a key modification: after myosin immobilization on the surface, MAVA was added to the chamber in the absence of ATP and incubated for 2 minutes, before actin was introduced. This allowed Mava to interact with myosin in a nucleotide-free, pre-actin binding state. Subsequently, the assay proceeded identically to **Setup 1,** with actin addition, motility initiation, and drug washout. Using these two configurations, we compared the effects of MAVA on full-length PsoM-II and PsoS1.

**Fig. 7.**
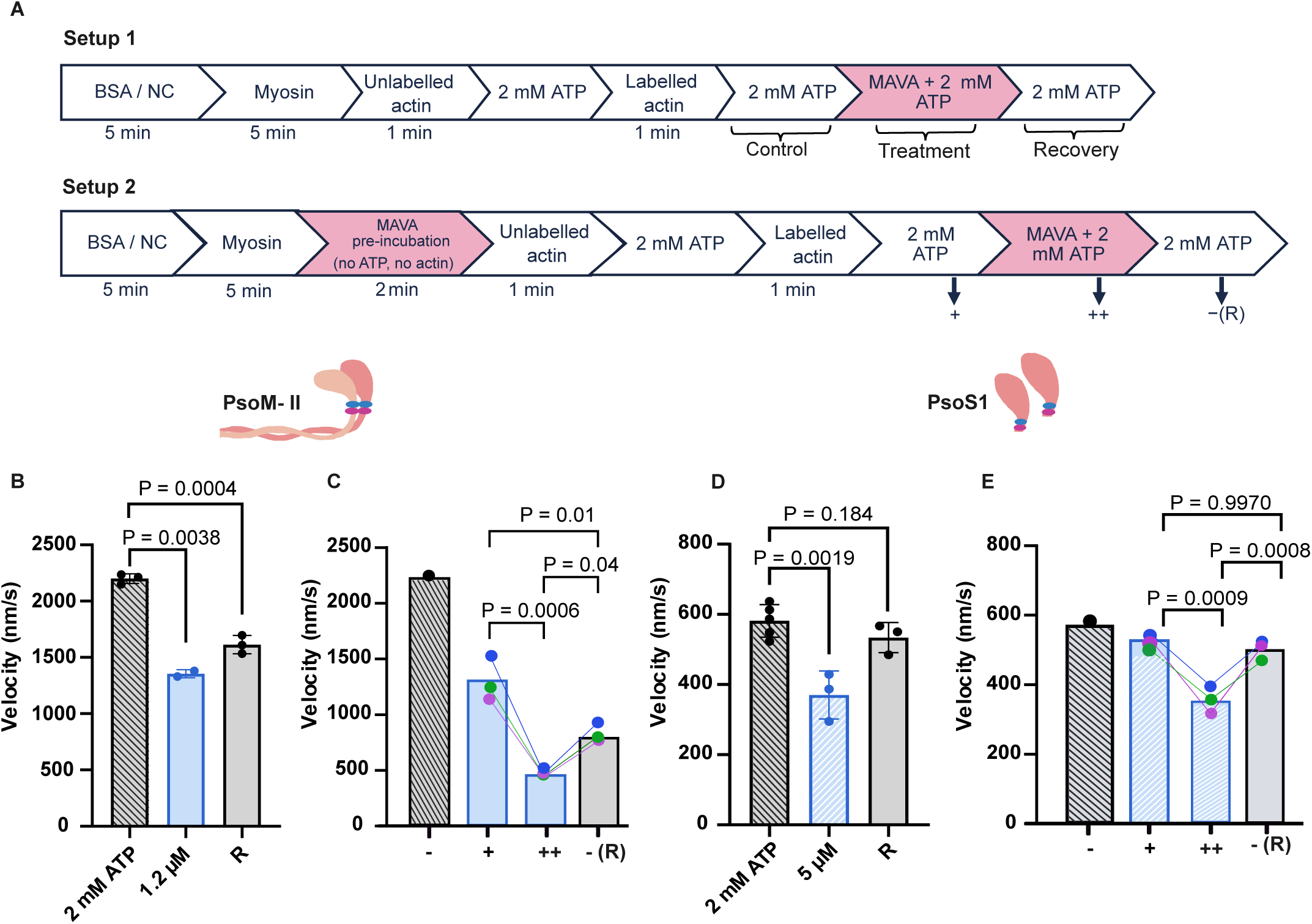
Effects of MAVA on full-length PsoM-II and single-headed PsoS1. **(A)** Schematic representation of the standard in vitro motility assay (Setup 1) and modified assay (Setup 2). In **setup 1**, myosin is immobilised on BSA coated surface. After addition of F-actin and 2 mM ATP motility is recorded (control). MAVA is added to the same chamber and myosin inhibition is recorded (treatment). Lastly, MAVA is washed off and motility is recorded at 2 mM ATP (recovery). In **setup 2**, MAVA is added to immobilized myosin in the absence of ATP. The later steps are same as in setup 1. **(B)** Effect of 1.2 µM MAVA on full-length PsoM-II using Setup 1. Actin-filament sliding velocity was recorded under control conditions (2 mM ATP), after addition of 1.2 µM MAVA, and followed by recovery after MAVA washout (R). **(C)** Motility conditions as described in Setup 2. The first bar (**−**) represents motility at 2 mM ATP (control), the second bar (**+**) represents motility following pre-incubation with 1.2 µM MAVA, the third bar (**++**) represents motility at 1.2 µM MAVA and 2 mM ATP, and −(R) represents recovery after MAVA washout. Note that in setup 2, the motility at 2 mM ATP (1^st^ bar, control) was conducted in a separate parallel chamber under identical experimental conditions of surface myosin density, 2 mM ATP and 22 °C. **(D and E)** Effect of 5 µM MAVA on PsoS1 using Setup 1 **(D)** and Setup 2 **(E)**. Experiments were performed at 22 °C, unless otherwise stated. Connected data points of the same color represent measurements obtained from the same motility chamber across successive conditions. N = 2-5, n = 100-150. Error bar: mean ± SD. Statistical significance was assessed using an unpaired two-tailed Student’s t-test (setup 1) and ANOVA (setup 2). N = number of chambers, n = number of filaments.

For PsoM-II, treatment with 1.2 µM MAVA using **Setup 1** resulted in a reduction of ∼39% in actin sliding velocity (Fig. 7B) and the partial activity was restored (∼73 %) after Mava washout. The measurements with **Setup 2** revealed overall stronger inhibition of myosin activity with reduction in actin gliding speed by about 40% (Fig. 7C) after pre-incubation of myosin with Mava. Subsequent addition of Mava to the motility buffer reduced the speed by ∼80 % of the initial non-drug condition, and led to only partial recovery (to about 38 % of the control condition) after drug washout. This recovery was less than in setup 1.

For PsoS1, 1.2 µM MAVA did not produce a significant effect on actin gliding speed. Therefore, experiments were performed at a higher concentration of 5 µM MAVA. As shown in Fig. 7D, under setup 1 conditions, a significant reduction in actin sliding velocity of ∼35% was observed in the presence of 5 µM MAVA, that recovered completely after drug removal (P = 0.18). In setup 2 (Fig. 7E), the actin gliding velocities remained unperturbed when the MAVA was preincubated with myosin prior to addition of actin. MAVA addition together with 2 mM ATP led to ∼40% reduction in velocity and reached full recovery post MAVA washout from the flow cell unlike for full-length PsoM-II in Fig. 7C. Nearly complete recovery indicates fully reversible MAVA binding for PsoS1.

To further investigate the dominant inhibitory effect of mavacamten requiring a dimeric myosin form, similar experiments were performed using psoas heavy meromyosin (Pso-HMM) to understand if subfragment-2 (S2) region of the MyHC is sufficient for this inhibitory effect. Pso-HMM was prepared by digesting PsoM-II with chymotrypsin (Fig. S5). We determined whether dimeric Pso-HMM behaves more similar to single-headed PsoS1 or full length PsoM-II (Fig. S6). In setup 1, the velocity was reduced by about 27 % at 5 µM MAVA, and full recovery was observed post drug-removal. In setup 2, preincubation of HMM with MAVA caused 30 % reduction in speed, which reduced further with subsequent addition of MAVA with 2 mM ATP to 52 %. The myosin activity recovered only partially to about 65 % after the treatment. Interestingly, the inhibitory and recovery profiles of Pso-HMM resembled PsoM-II rather than PsoS1 (Fig. 7), suggesting that the presence of two myosin heads is important to reproduce the dominant inhibitory effect of MAVA. However, dimeric HMM was less sensitive when compared with full-length myosin, which exhibited about 74 % reduction in velocity and incomplete recovery to 58 % at 5 µM MAVA (Fig. 3C).

We further evaluated whether the inhibitory effect of MAVA on pre-actin-bound and post-actin-bound myosin configuration is conserved across different myosin isoforms. Similar experiments were performed using native SolM-II, βM-II, and DiaM-II (Fig. S7, Fig. S8, and Fig. S9). Near IC_50_ values of MAVA concentrations were employed in these assays for respective myosin isoforms, for e.g., 100 nM for SolM-II, 250 nM for βM-II, and 1 µM for DiaM-II. Consistent with the observations obtained for PsoM-II, pre-incubation of myosin with mavacamten in the absence of actin resulted in a reduction in actin filament sliding, which decreased further when MAVA was added in final motility buffer. Recovery following drug washout was also observed across all tested myosin types. Interestingly, as shown earlier (Fig. 7C), for all the myosin isoforms tested, the reduction in velocity associated with actin-bound myosin configuration was largely reversible upon MAVA washout, but did not reach full recovery.

## Discussion

The central finding of this study is that MAVA-mediated inhibition of myosin II motor function is not limited to β-cardiac myosin in the heart but also influences diaphragm myosin, which contains a mixture of fast and slow skeletal myosin II isoforms. Our studies show that the extent of inhibition varies among isoforms and depends on myosin heavy- and light-chain composition, two-headed structural architecture, and thin-filament regulation. Moreover, recovery kinetics after drug removal differed among distinct tissue-derived myosins. For diaphragm, M. soleus, M. psoas, and β-cardiac myosin preparations, MAVA sensitivity follows a clear isoform-dependent hierarchy, with slow skeletal and β-cardiac myosins more sensitive than fast skeletal myosin, and diaphragm myosin showing an intermediate response, i.e., **SolM-II > βM-II > DiaM-II > PsoM-II**. Mechanistically, comparing full-length myosin, HMM, and S1 indicates that the dominant inhibitory effect is strongly enhanced when the two-headed motor and proximal S2 are retained, supporting a model in which MAVA reduces the number of actin-accessible myosin heads. Furthermore, MAVA directly affects the kinetics of the ATP-dependent actomyosin cross-bridge cycle. Thus, we distinguished two modes of action of MAVA underlying its inhibitory function: MAVA binding to myosin in two different conformations, 1) when myosin is not interacting with actin and 2) when myosin is undergoing the active cross-bridge cycle (Fig. 8), both contributing to inhibition and reduction of actin gliding speed.

**Fig. 8.**
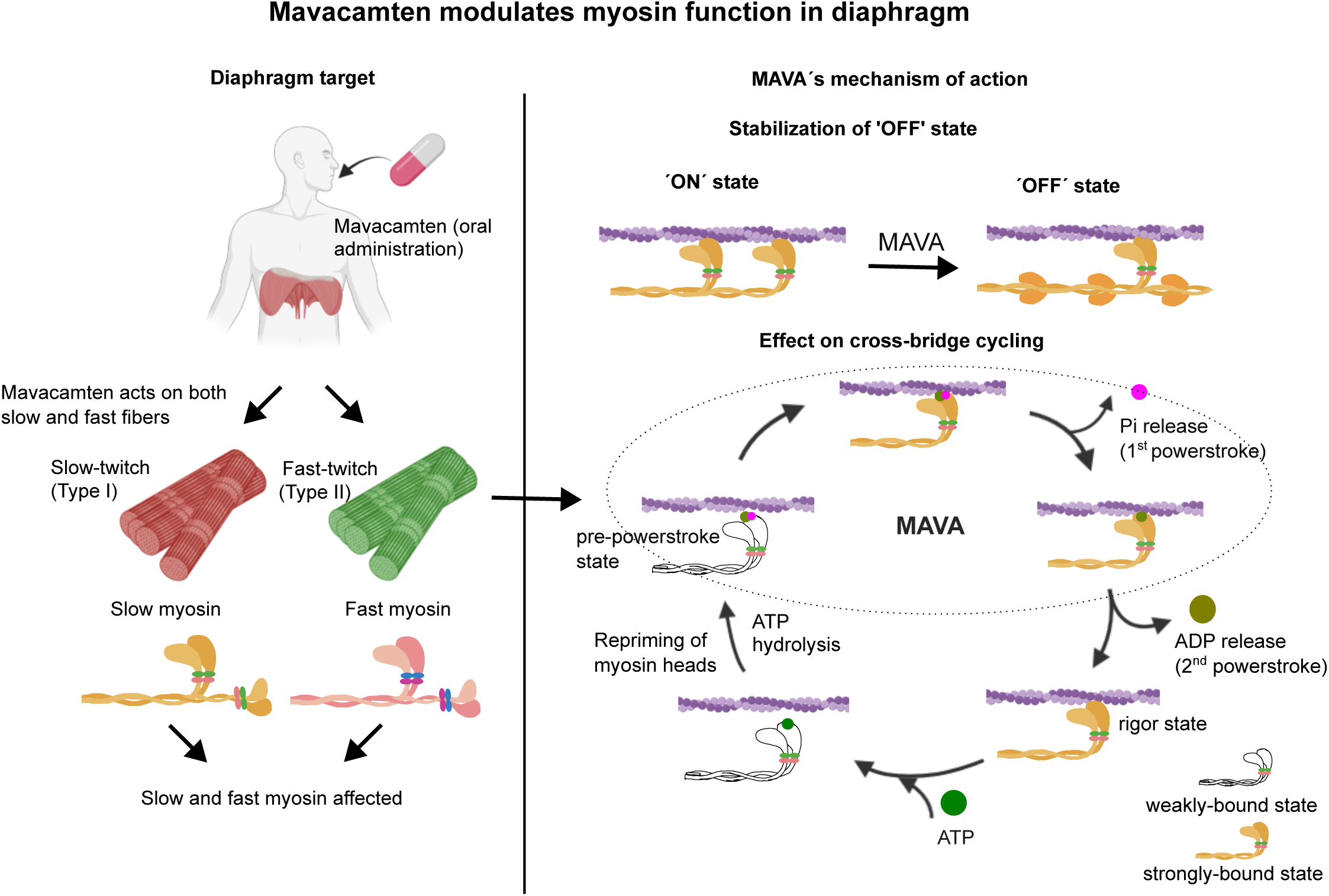
Proposed model of MAVA action on diaphragm myosin. **Left panel-**MAVA acts on both slow and fast fibres of the diaphragm following oral administration. The drug targets myosin at the molecular level and affects both slow and fast myosin isoforms. **Right panel-**MAVA likely stabilizes the ‘OFF’ myosin states, thereby reducing the number of active myosin heads available for actin interaction and cross-bridge cycling. Simplified actomyosin crossbridge cycle with key transition states are shown to indicate the states affected by MAVA exposure based on previous studies and our observations, indicated in dotted oval, which includes transitions from weak-strongly bound acto-myosin states, and Pi) release.

### Isoform-dependent sensitivity of myosin to MAVA

SolM-II has shown the highest sensitivity to MAVA in our experiments. The observed strong effect of MAVA on SolM-II is mechanistically plausible as they share the same MyHC. However, the two-fold difference in the IC_50_ values of SolM-II and βM-II was surprising and suggests that myosin heavy chains alone do not fully explain MAVA sensitivity. In our earlier studies, we showed that the light chains contribute to the mechanical and kinetic differences between myosin isoforms (*38*, *39*). SolM-II contains an additional myosin LC isoform, MLC1sa (along with MLC1sb/v), which is longer and more positively charged. MLC1sa decreased actin and thin-filament gliding on β-myosin molecules by nearly 3-fold, owing to the same extent of decreased maximum actomyosin detachment rates (*38*, *39*, *44*). We propose that MLC1sa in the SolM-II confers an additional MAVA-induced inhibitory effect, possibly by affecting the myosin conformations or the kinetics of ATPase cycling, or both. These studies further reinforced the concept that light chains are not passive structural elements but active modulators of myosin kinetics and mechanics (*45*). Several studies have suggested that the long N-terminal extension containing approximately 40 amino acid residues enriched in lysine and proline residues of the ELC can interact directly with actin (*46*–*49*). Although still debated, it has been suggested that the positively charged ELC residues may weakly interact with the negatively charged C-terminal region of actin, modulating actomyosin cross-bridge kinetics by reducing the actomyosin detachment rate (*50*). In addition, a slower ADP release rate from actomyosin, through its allosteric effect has also been proposed as a mechanism for reduced detachment rate, consequently decreasing filament-sliding velocity (*39*). These may explain SolM-II’s greater susceptibility to MAVA inhibition.

The low sensitivity of PsoM-II further supports isoform-dependent drug action, suggesting that MAVA preferentially affects myosin states in slow/βM-II. A direct molecular explanation for the weak responsiveness of fast skeletal myosin is provided by the structural work (*32*). In the MAVA-bound β-cardiac myosin structure, the drug binds in a pocket between the N-terminal/U50 region and the converter region. Importantly, the class-II myosin sequence alignment identified two drug-contact residues that differ between cardiac and fast skeletal myosin: β-cardiac MyHC Tyr164 is replaced by Phe and Asn711 by Ser in fast skeletal myosin (*32*). These substitutions were linked to the approximately ten-fold weaker MAVA affinity of fast skeletal myosin. Consistent with this structure-based analysis, we observed nearly a 10-fold difference in the IC_50_ of PsoM-II and βM-II in functional measurements. While the sequence differences are a strong molecular candidate, further differences between these two myosin isoforms include light-chain composition, actomyosin cross-bridge kinetics, and their equilibrium between active and inhibited conformations, which are expected to contribute to the overall functional effects (*45*, *51*).

The intermediate DiaM-II IC_50_ is consistent with the combined contribution of slow and fast myosin populations in the mixed diaphragm preparation. When measured with pure slow (SolM-II) and fast (PsoM-II) myosin isoforms, fast myosin requires nearly 20-fold higher MAVA to reach the IC_50_ than slow myosin. In a mixed population, slow myosin, inhibited at much lower MAVA concentrations, may exert a drag force that slows overall fast myosin-driven actin filament gliding. Mechanistically, the difference between slow and fast myosin sensitivity may also relate to the ability of different isoforms to populate inhibited states. MAVA has been shown to stabilize folded-back or autoinhibited myosin conformations (*27*). Structural inhibition or folded back states of subpopulation of motors are less likely to reduce gliding speed unless the myosin density ready to bind actin filaments is reduced below a threshold that cannot support the smooth motility or when they can weakly interact with actin and generate drag force on the moving filaments driven by the uninhibited myosin population. Therefore, we interpret the observed effects as primarily due to MAVA’s effect on actomyosin attachment/detachment kinetics, with slow and β-cardiac myosins more strongly inhibited. We further observed that dimeric myosins are more sensitive than monomeric motor heads and that full-length myosin is the most sensitive.

### Reversibility of MAVA inhibition and myosin isoform-dependent recovery after drug washout

After removing MAVA from the motility chamber, myosin activity recovered in DiaM-II, PsoM-II, SolM-II, and βM-II, either partially or fully, supporting MAVA as a reversible allosteric inhibitor of myosin.

Previously, it is shown that MAVA affects the actomyosin ATPase cycle by slowing phosphate release and reducing the number of myosin heads that productively enters force-generating actomyosin states. By sequestration of myosin heads, MAVA directly impacts actomyosin attachment kinetics. Reduced number of actomyosin interactions may thus slow the actin filament gliding velocity. Presumably, drug release from myosin allows it to progress through the normal ATPase cycle. However, recovery post-drug washout was not identical among all isoforms, likely due to differences in the affinity of the drug for myosin isoforms. Interestingly, contrary to expectations, DiaM-II and PsoM-II showed better recovery at higher MAVA concentrations. One possible explanation is that higher MAVA concentrations may orient myosin heads in a more ordered inhibited conformation during drug exposure and upon washout, these heads assume a more favourable orientation for actin interaction, leading to improved recovery of motility. This interpretation may be especially relevant for fast myosin, which is less sensitive to MAVA. Thus, lower MAVA concentrations may inhibit only a subset of myosin heads, producing a more heterogeneous mixture of active or weakly inhibited heads. After washout, this heterogeneity result in incomplete or variable recovery.

DiaM-II with added complexity due to a mixture of slow and fast myosin isoforms, at higher MAVA concentrations, both slow and fast myosins may be more uniformly shifted into inhibited states, allowing a more coordinated return of actin gliding after washout. The apparently improved recovery at selected intermediate MAVA concentrations cannot be interpreted as a simple concentration-dependent effect, because recovery was again limited at 100 µM, where motility is strongly inhibited. Thus, the explanation that higher drug concentrations generate a more uniform reactivated population remains plausible for some concentrations, beyond which myosin appears to enter a ‘deeply inhibited state’ that cannot fully recover, consistent with earlier observations in myofibril-level studies (*52*).

SolM-II showed incomplete recovery across the tested MAVA concentrations, consistent with its high sensitivity and presumably high affinity. Our observations are consistent with a previous report showing slower force redevelopment after longer MAVA exposure in cardiac myofibrils (*52*). In our DiaM-II experiments, extending the post-washout period did not further improve recovery (Fig. S4), arguing against insufficient wash duration and incomplete removal of residual MAVA in solution and suggesting that some myosin heads assume inhibited states that did not recover over the tested period.

Overall, the recovery data show that MAVA inhibition is reversible, but the extent of recovery depends on myosin isoform and MAVA concentration.

### Mechanism of MAVA-mediated inhibition

We observed a significant difference in MAVA’s effect on full-length myosin (PsoM-II) compared to monomeric myosin (PsoS1), with the dimeric form being more sensitive. PsoS1, which lacks the dimeric architecture necessary for forming interacting-head motif (IHM)-like head sequestration, therefore reveals the direct effect of MAVA on the chemomechanical cycle. In full-length PsoM-II, MAVA strongly reduced actin filament sliding velocity. At 1.2 µM MAVA, while full-length PsoM-II showed around 37% inhibition (Fig. 6B), PsoS1 showed a modest reduction of ∼ 17% (Fig. 6C). This difference indicates that the direct effect on the actomyosin crossbridge cycle alone is insufficient to explain the inhibitory effect of MAVA in full-length myosin. Similar differences were observed earlier, with MAVA promoting the super-relaxed (SRX) or low ATP turnover state that is more pronounced in dimeric motors than in single myosin heads (*27*, *28*).

Recent cryo-EM studies provide direct structural evidence supporting the existence of sequestered myosin head structures at high concentrations of MAVA of 20 -to 50 µM. These structures were observed in intact myofibrils, thick filaments, and at the individual molecule level (*10*, *14*). However, MAVA induced promotion of these states was less pronounced in individual myosin molecules with a modest increase (up to 20%) in the number of IHM-like sequestered head states. In dimeric myosin, MAVA stabilizes the ADP·Pi-bound pre-powerstroke state, with a restricted lever-arm configuration and actin-binding-cleft dynamics, along with slow transition toward force-generating states (*10*, *32*). Important to note that these studies primarily indicate that the Pi release from the pre-powerstroke states is slower. Our recent optical trapping-based single-molecule function investigation provides direct evidence of delayed powerstroke in the presence of MAVA, which may be associated with delayed Pi release after the transition from the weak to the strongly bound actomyosin state (*53*). This suggests that, beyond slowing the weak-to-strong actomyosin binding transition, MAVA further delays power stroke generation, which typically occurs in less than 2 ms after strong actomyosin binding.

Pso-HMM, which retains two heads together with approximately 40 heptads of proximal S2, resembled full-length PsoM-II than PsoS1 for its response to MAVA (Fig. S6). However, the inhibition in Psoas HMM was less pronounced than in PsoM-II, indicating that distal tail region is required for the full-length-like response under our assay conditions.

### Inhibitory modes of action of MAVA

We found that preincubating myosin with MAVA in the absence of actin filaments and ATP was sufficient to alter myosin activity, resulting in a reduction of actin motility by about 40%. Subsequent addition of MAVA and 2 mM ATP in the same chamber, further decreased actin velocity, with a combined inhibition of approximately 80%, which was greater than when MAVA was added after actomyosin complex formation (Fig. 7B-C).

Therefore, we assume that MAVA addition after actomyosin complex formation reveals the drug’s effect on actomyosin chemomechanical cycle (Fig. 7B), resulting in the delayed power stroke generation and altered powerstroke size, as suggested by our recent findings (*53*) in combination with slower Pi release and transition from weak-to strongly-bound AM states (*31*). Pre-incubating myosin with MAVA must be a different MAVA-induced conformation of myosin that affects the actin filament gliding.

Interestingly, our recovery data shows that myosin function inhibition during cross-bridge cycling appears readily reversible after drug washout than myosin inhibition in the pre-inhibited step (before actin addition) (Fig. S6, S7,S8 and S9), supporting the idea that MAVA-bound myosin states differ in reversibility on the time scale of our assay, likely due to myosin populations in different MAVA-bound states. During active cycling, removal of the drug may allow a myosin population to re-enter the normal ATPase cycle, supporting motility. On the other hand, MAVA may shift a fraction of myosin into a possible ‘deeply inhibited’ state or a slowly reversible inhibited state. It remains to be examined whether ‘deeply inhibited’ states’ represent the sequestered myosin heads.

Collectively, these data support a distinct state-dependent response whose magnitude depends on both myosin isoform and molecular architecture, i.e., single-headed *vs* dimeric motor.

### Native thin filaments and sensitivity to MAVA

Under activating conditions at pCa 4.0, DiaM-II showed 2-fold greater sensitivity to MAVA in the presence of native thin filaments than in unregulated F-actin (IC_50_ of 1352 nM, and 691 nM for F-actin and nTF, respectively), suggesting that nTF regulatory proteins modify M-II’s functional response to MAVA. One possible explanation for the lower IC_50_ observed with nTFs may be that MAVA reduces the number of myosin heads available for strong actin binding. In a bare F-actin system, reduced myosin density directly reduces actin sliding velocity (*54*). In a regulated native thin filament system, apart from Ca^2+-^induced access of actin to myosin by relieving Tm-Tn-mediated steric blocking, strong myosin binding to actin also helps maintain the thin filament in an ‘open’ state, allowing myosin to readily associate with actin filaments. Furthermore, Tm-Tn components increase Pi and ADP release rates (*55*, *56*) from myosin, increasing ATP turnover rates and, subsequently, TF gliding speed. Therefore, if MAVA decreases the number of available or strongly bound myosin heads, it may also reduce cooperative thin filament activation. Thin filament regulation involves controlled access of myosin to actin and that strong myosin binding itself feeds back to activate the filament (*57*, *58*). Furthermore, if nTFs regulate Pi-release rates, then MAVA-mediated slowing of Pi release may have a larger functional effect with nTFs. The importance of the native thin-filament context is also supported by our recent work showing that tissue-specific nTFs can alter both gliding speed and Ca^2+^ sensitivity (*44*). This provides independent support that the TFs are an active determinant of ensemble myosin function.

Accordingly, increased MAVA sensitivity in diaphragm nTFs is likely to be an emergent property of the regulated actomyosin system rather than indicative of MAVA directly targeting troponin or tropomyosin function.

### Physiological implications for diaphragm function under MAVA treatment

As the diaphragm is the principal inspiratory muscle, any pharmacological modulation of diaphragm myosin has direct physiological consequence. The mixed muscle fiber type composition allows this muscle to recruit the specific fiber types. Here, we show that DiaM-II is clearly sensitive to MAVA, s and that MAVA may affect diaphragm myosin activity in patients receiving MAVA for hypertrophic cardiomyopathy.

The diaphragm exhibits remarkable plasticity under pathological conditions. For example, in severe chronic obstructive pulmonary disease (COPD), human diaphragm fibers shift toward slow-twitch phenotypes, with reduced force generation, decreased MyHC content, impaired Ca^2+^ sensitivity, and slower cross-bridge kinetics, suggesting adaptive remodeling with impaired function (*59*–*62*). Recent work showed that mechanical ventilation induces rapid disuse-related atrophy, with diaphragm fibers from critically ill patients showing severe contractile weakness and activation of proteolytic pathways within days (*63*, *64*). Notably, diaphragm myosin can be trapped in an energy-sparing super-relaxed state, linked to respiratory muscle “hibernation” and reduced diaphragm contractile function in mechanically ventilated ICU patients (*65*).

MAVA is therapeutically beneficial in treating HOCM as excessive sarcomere activity contributes to hypercontractility and left ventricular outflow tract obstruction (*19*). However, although MAVA may be well tolerated in a healthy diaphragm, it is conceivable that if a patient receiving MAVA already has under-functioning diaphragm, for example, patients with chronic respiratory disease, ICU-acquired weakness, prolonged mechanical ventilation, neuromuscular disease, etc., further stabilization of inhibited myosin states could potentially reduce the diaphragm’s ability to function normally. So far, this is not reported clinically for MAVA, but it is a very valid concern arising from our data.

Rat diaphragm shows an age-related increase in the proportion of slow MyHC (*66*). Slow fibers are important for fatigue-resistant, repetitive respiratory activity (*67*); preferential inhibition of slow myosin could theoretically influence endurance of diaphragm muscle. As MAVA strongly inhibits slow myosin, and therefore Type-I muscle fibers, fast fibers may be more readily recruited to compensate for the loss of function, even under resting conditions. Importantly, with a total plasma concentration range of MAVA in a treated patient estimated at about 1300-2600 nM (*29*), this concentration is sufficiently high to influence both slow and fast fibers. Although the free drug concentration may be lower, the effects on slow fibers can be expected. Our recovery data provides additional insight. For diaphragm physiology, contractile recovery may depend on the kinetics of reactivation of inhibited myosins, not simply on drug clearance. This may be particularly relevant during fluctuating respiratory demand, where rapid recruitment of additional force-generating myosin heads may be required.

Clinically, MAVA has been evaluated primarily as a cardiac myosin inhibitor for symptomatic HOCM (*19*). There is currently no strong clinical evidence that MAVA directly causes diaphragm dysfunction in treated HCM patients. MAVA inhibited slow-twitch skeletal muscle myosin and weakly inhibited fast-twitch skeletal myosin in vitro, but skeletal muscle function was not affected in nonclinical *in vivo* safety studies, assessed by grip strength, locomotor function, or respiratory function in dogs (*68*). Clinical HCM trials report that MAVA improves symptoms by reducing cardiac hypercontractility and outflow obstruction and long-term clinical studies have reported sustained improvement in cardiac function without major safety concerns (*69*). Therefore, in most patients, the net clinical effect is considered beneficial as cardiac unloading and improved hemodynamics may reduce dyspnea. However, from a skeletal muscle and diaphragm physiology perspective, our data suggest that skeletal and respiratory muscle effects may occur in vulnerable subgroups, at high exposure, or in patients with pre-existing diaphragm or respiratory dysfunctions.

*Conclusion.* Collectively, we find that the diaphragm, comprising slow and fast skeletal muscle fibers, is sensitive to MAVA treatment. Since the function of Type-I slow-twitch muscle fibers is even more strongly inhibited than the β-cardiac myosin and most muscles contain a mixture of Type-I and Type-II muscle fiber types, depending on the content of the Type-I fibers, various muscles likely experience the MAVA-mediated inhibition of function. Both MyHC and LC composition tune the probability of entering or remaining in these inhibited states, providing a mechanistic basis for the greater sensitivity of slow skeletal and β-cardiac myosins than fast skeletal myosin. This sensitivity is further expected to be modified by nTF regulation *in vivo* conditions. This fundamental study is highly clinically relevant, providing a framework for future investigations into both the short- and long-term effects of the MAVA-based treatments on respiratory function, including the side effects of myotropes for cardiac conditions.

## Limitations of the study

Our studies show that diaphragm and slow myosin isoforms are sensitive to MAVA under controlled *in vitro* conditions. These *in vitro* observations will further benefit from the direct measurement of additional parameters, such as force generation and sarcomere contractility, in muscle fibers and/or intact diaphragm strips. Animal models with short and long-term exposure to MAVA combined with respiratory function monitoring will provide key information. Finally, for direct clinical relevance to humans, clinical observations of diaphragm function and studies of the human diaphragm would be beneficial, as species-specific differences and disease state may modify MAVA sensitivity and its collateral effects *in vivo*.

## Materials and Methods

### Muscle tissues

Muscle tissues were obtained from 6 months old New Zealand White rabbits (Crl:KBL, NZW). Animals were euthanized in compliance with the German Animal Protection Act (§7, sacrifice for scientific purposes). Shared tissues were used from animals approved under authorization number 18A255. Rabbits registered under reference number G43290 were supplied by Charles River France. All experimental procedures adhered to the regulations and guidelines of the Lower Saxony State Office for Consumer Protection and Food Safety and Hannover Medical School, Germany.

### Native myosin II

Full-length diaphragm myosin II, fast skeletal myosin II (PsoM-II) and slow skeletal myosin II (SolM-II), β-cardiac myosin were purified from rabbit diaphragm, *M. psoas* and *M. soleus* muscle, and ventricle respectively, using a high-salt extraction buffer containing 0.5 M NaCl, 10 mM HEPES (pH 7.0), 5 mM MgCl₂, 2.5 mM MgATP, supplemented with fresh 2 mM DTT and 0.5 mM AEBSF. Myosins were extracted from 100-150 mg of diaphragm, *M. soleus, M. psoas,* and left ventricular tissue that was first pulverized using a liquid nitrogen–cooled mortar and pestle. The resulting powder was incubated in extraction buffer for 30 min on ice to isolate myosin. Cellular debris was removed by ultracentrifugation at 65,000 rpm (TLA 120.2 rotor; Beckman Coulter) for 1 h. The clarified supernatant was subsequently diluted 10-fold with ice-cold ultrapure water supplemented with 2 mM DTT and 0.5 mM AEBSF to induce myosin precipitation. Following incubation on ice for 40 min, the precipitated myosin was collected by centrifugation at 37,000 rpm (TLA110 rotor; Beckman Coulter) for 30 min. The resulting pellet was gently resuspended in myosin extraction buffer. Protein concentration was quantified using the Bradford protein assay. Purified myosin was snap-frozen in liquid nitrogen in the presence of 50% glycerol (small size ∼10 µl aliquots) to minimize freeze-induced damage and stored at −80 °C for long-term use. For the generation of single and double-headed myosin motors from native myosin II, full length psoas myosin II was digested with papain or chymotrypsin to generate single-headed sub-fragment 1 (myosin S1) or myosin HMM, respectively, as described earlier (*70*).

### Myosin heavy- and light-chain isoform SDS-PAGE

Distinct electrophoretic conditions were applied to resolve myosin heavy-chain (MyHC) isoforms. Skeletal muscle MyHC isoforms were separated using 6.5% acrylamide/bisacrylamide (AA/BisAA, 100:1) gels. The stacking gel consisted of 4.5% AA/BisAA supplemented with 30% glycerol, while the separating gel contained 8% AA/BisAA with 30% glycerol. 0.75 – 2.5 µg protein samples were loaded per lane. Gels (10 cm total length; 1.5 cm stacking and 8.5 cm separating) were run in a custom-made chamber at 4 °C temperature for 28.5 h, initially at 10 mA for 1 h, followed by 12 mA constant current. Proteins were visualized using Quick Coomassie stain (Biotrend, NB-45-00078-1L).

The protein composition of myosin light chains (LCs) and thin filament preparations were analyzed using 12.5% and 15% SDS–PAGE in custom-made gel chambers. Gels were prepared with an acrylamide/bisacrylamide ratio of 37.5:1 using RotiphoreseGel 30 (Roth, 3029.1). Approximately 2.5 -5 μg of protein was loaded per lane. Following electrophoresis, gels were stained with Quick Coomassie Stain (Biotrend, NB-45-00078-1L) for 30 min. Gel images were acquired using a Biostep Dark Hood (DH-30/32) imaging system with an exposure time of 200 ms.

### Cryosectioning and Immunofluorescence Staining

Frozen tissue samples from liq N2 or freshly excised tissues were embedded and cryosectioned using a cryostat maintained at −20 °C. Sections of 5 µm thickness were collected onto clean glass slides. Slides were air-dried and stored at 4°C until further processing. Prior to staining, sections were fixed with 4% paraformaldehyde (PFA) in PBS for 1 h at room temperature. After fixation, sections were washed twice with PBS (5 min per wash). Fixed cryosections were either processed immediately or stored in PBS at 4 °C and used within 2 days.

For permeabilization, cryosections were incubated in 0.5% Triton X-100 in PBS for 30 min at room temperature. Sections were then washed with PBS and incubated in blocking solution containing 5 % BSA in PBS for 30 min at room temperature to reduce nonspecific antibody binding. Primary antibodies were diluted in BSA blocking solution and applied to the sections for 1 h at room temperature. The following primary antibodies were used for fiber-type identification: mouse anti–slow myosin heavy chain (β-MyHC) (Sigma-Aldrich, M8421; 1:100 dilution) and rabbit anti–fast myosin heavy chain (Abcam, ab91506; 1:100 dilution). Following primary antibody incubation, sections were washed twice with PBS (5 min each). Secondary antibodies diluted in BSA were then applied for 1 h at room temperature. The following secondary antibodies were used: Alexa Fluor 555 donkey anti-mouse IgG (Thermo Fisher Scientific, A31570; 1:400 dilution) and Alexa Fluor 488 donkey anti-rabbit IgG (Thermo Fisher Scientific, A21206; 1:400 dilution).

After secondary antibody incubation, sections were washed with PBS (5 min). FITC-WGA (wheat germ agglutinin) at 1:100 dilution -was added to sections for 2 minutes at room temperature to mark the membrane of muscle fibers followed by a 5 min wash with PBS. Nuclei were stained with DAPI (1:12500 dilution in PBS) for 2 min at room temperature, followed by a final 5 min PBS wash. Sections were briefly rinsed with distilled water, partially air-dried, and mounted using Fluoroshield mounting medium. Coverslips were applied and sealed with a sealant. Slides were stored at 4 °C until imaging.

### Actin filament polymerization

G-Actin was isolated from chicken *M. pectoralis major* as previously described (*71*). For in vitro motility assays, actin filaments (F-actin) were polymerized by incubating G-actin in polymerization buffer (p-buffer) containing 5 mM Na-phosphate, 50 mM K-acetate, and 2 mM Mg-acetate, supplemented with 0.5 mM AEBSF protease inhibitor, overnight at 4 °C. Fluorescent filaments were generated by adding an equimolar amount of rhodamine-phalloidin (Sigma-Aldrich). Unlabelled filaments were prepared the same way but without rhodamine-phalloidin.

### Native Thin Filaments

Native thin filaments (nTF) from rabbit diaphragm were isolated using a recently published protocol for small tissue samples (*44*). Briefly, tissue samples (50–100 mg) were mechanically disrupted in liquid nitrogen using a pre-chilled mortar and pestle. The powdered tissue was transferred into skinning solution containing 3 mM Na_2_ATP, 5 mM Mg-acetate, 5 mM ethylene glycol-bis (2-aminoethylether)-N,N,N′,N′-tetraacetic acid (EGTA) (Sigma-Aldrich, E4378), 50 mM creatine phosphate (Sigma-Aldrich 2380), 5 mM KH_2_PO_4_, 8 mM NaN_3_, supplemented with fresh 1 mM DTT (Sigma-Aldrich, D0632-10G), 1× cOmplete EDTA-free Protease Inhibitor Cocktail (Roche, 05056489001), 20 mM BDM (Sigma, B0753), and 1% Triton X-100 (Roche, 11332481001) and incubated on ice. After removal of the skinning solution by low-speed centrifugation, pellets were washed sequentially in washing buffer (same as skinning solution, but without BDM and Triton X-100) to eliminate detergent and BDM, followed by resuspension in homogenization buffer containing 10 mM imidazole, 25 mM propionic acid (Sigma-Aldrich, P1386), 10 mM caffeine (Sigma-Aldrich, C0750), 10 mM creatine phosphate, 2 mM MgCl_2_, 3 mM EGTA, 2 mM MgATP, and freshly added 2 mM DTT and 1 mM 4-(2-Aminoethyl) benzene-1-sulfonyl fluoride (AEBSF) Hydrochloride (AppliChem, A1421). Homogenization was performed using a Precellys system with ceramic beads (two 2.8 mm beads and five 1.4 mm beads per tube), and myofibrils were collected by centrifugation. The pellet was subsequently resuspended in suspension buffer (same as homogenization buffer with added 10 mM MgATP and 10 mM DTT) and subjected to Dounce homogenization to separate thin filaments from supernatant after centrifugation steps. nTFs were pelleted by ultracentrifugation (TLA 120.2 rotor). The pellet was resuspended in assay buffer containing 25 mM imidazole-HCl, pH 7.2, 25 mM NaCl, 4 mM MgCl₂, 1 mM EGTA, supplemented with 2 mM DTT and 1 mM AEBSF. Thin filaments were fluorescently labeled with rhodamine-phalloidin (0.25 nM) and further purified by running nTF sample over a sucrose cushion with a centrifugation at 70,000 rpm for 30 min. The final pellet was gently resuspended in assay buffer prior to use. Protein concentrations were determined using the Bradford assay. The thin filaments were stored at 4°C and used within 2 days in the motility assays.

### *In vitro* motility assay

In vitro motility assays were performed using full-length myosins immobilized on BSA-coated flow cell surfaces. HMM and S1 were immobilized on nitrocellulose-coated surface. We followed the motility protocol as described earlier with some modifications (*72*). Assay buffer containing 1 mg/ml BSA (25 mM imidazole-HCl, pH 7.2, 25 mM NaCl, 4 mM MgCl₂, 1 mM EGTA, and 2 mM DTT) was introduced into the flow cell and incubated for 5 min, after which myosin was added and allowed to bind for an additional 5 min. Unbound myosin was removed by washing with extraction buffer. Non-functional myosin heads were blocked by introducing 0.25 μM short, unlabeled F-actin for 1 min, followed by the addition of 2 mM ATP to release bound actin and expose active myosin motor heads. ATP was removed by washing with assay buffer. Rhodamine phalloidin-labeled F-actin was then introduced for 1 min, and excess filaments were removed by washing. Motility was initiated by addition of assay buffer containing 2 mM MgATP and an oxygen-scavenging system (18 μg/ml catalase, 0.1 mg/ml glucose oxidase, 10 mg/ml D-glucose, and 10 mM DTT). To assess drug dependence, motility measurements were conducted at drug concentrations ranging from 1 nM -100 µM Mava. For measuring the recovery, following drug treatment, the flow cell was thoroughly washed by using 3 chamber volumes 2 mM MgATP solution.

### Motility assay with native thin filaments

Motility assays with native thin filaments (nTF) followed the similar procedure as described above for F actin motility, with rhodamine-phalloidin-labelled nTF introduced in place of labelled F-actin. Assay and motility buffers contained 1 mM EGTA, and free Ca^2+^ was set by adding CaCl_2_ to reach pCa 8.0 (low Ca²⁺) or pCa 4.0 (activating). The total CaCl₂ required for each pCa was calculated with PyChelator (*73*). Ca^2+^-mediated regulation of each nTF preparation was verified by comparing motility at pCa 8.0 (low Ca^2+^ condition) and pCa 4.0 (high Ca^2+^ condition), before MAVA was tested. MAVA concentration–response experiments with nTF were performed at pCa 4.0. nTF were stored at 4 °C and used within 2 days, to limit loss of tropomyosin–troponin from the filaments. Image acquisition and velocity analysis were followed as described for F-actin.

### Total Internal Reflection Fluorescence Microscopy

In vitro motility assays were imaged using an inverted microscope equipped with a custom-built four-channel simultaneous-imaging total internal reflection fluorescence (TIRF) system, as described previously (*72*). Rhodamine-phalloidin–labeled actin filaments were excited with a 532 nm Nd:YAG laser (Coherent, Compass 315M-150 SL). Fluorescence emission was detected using an Andor camera (Andor Technology, iXon DV887). The stage temperature was maintained at approximately 22 °C throughout image acquisition. Time-lapse image sequences were recorded at a temporal resolution of 0.2 s per frame (5 frames s⁻¹). For each measurement, movies consisting of 300 consecutive frames were acquired using Andor Solis software, generating 256 × 256 pixel 32-bit TIFF stacks with a spatial sampling of 156 nm per pixel.

### Data Analysis and Statistics

Image stacks were analysed in Fiji, an open-source platform for biological image analysis, and smoothly moving filaments were manually tracked using the MTrackJ plugin in Fiji v1.54g (*74*, *75*). Filament-velocity distributions for each experimental condition were fitted with a Gaussian function, from which the mean ± SD was obtained. GraphPad Prism v.9.5.1 was used for data plotting, nonlinear regression fits, and statistical analyses.

MAVA concentration–response curves were fitted by nonlinear regression using a four-parameter logistic model with a variable slope to determine the half-maximal inhibitory concentration (IC_50_). For motility experiments involving comparisons between two experimental conditions, statistical significance was assessed using an unpaired two-tailed Student’s t-test. For the modified state-dependent experiments, in which multiple experimental conditions were compared, statistical significance was assessed using analysis of variance (ANOVA). Data are presented as mean ± SD unless stated otherwise. Statistical tests are mentioned in the respective figure legends, wherever applicable.

To estimate the percentage of slow and fast fibres, the fluorescent images from 28 different cryosections (costal region) of diaphragm tissue were manually analysed. Statistical significance was assessed using unpaired two-tailed Student’s t-test.

## Data availability

All the data supporting the findings of this study are available within the article and the supplementary information files.

## General

We are grateful to Petra Uta for the technical assistance in protein preparations. We thank Tianbang Wang and Emrulla Spahiu for critical comments on the manuscript.

## Author contributions

MA conceived the project. MA and AS designed the experiments with input from AN and TK. AS performed all the experiments and analyzed the data. AS and TH performed immunofluorescence assay. MA and AS interpreted the data. MA supervised the project. AS and MA wrote and edited the manuscript. All authors contributed to editing the manuscript.

