## Supplemental Information for "β-Myosin inhibition in skeletal muscle: Mavacamten’ s impact on the diaphragm"

This Supplementary Information file includes,

- Supplementary figures 1-9

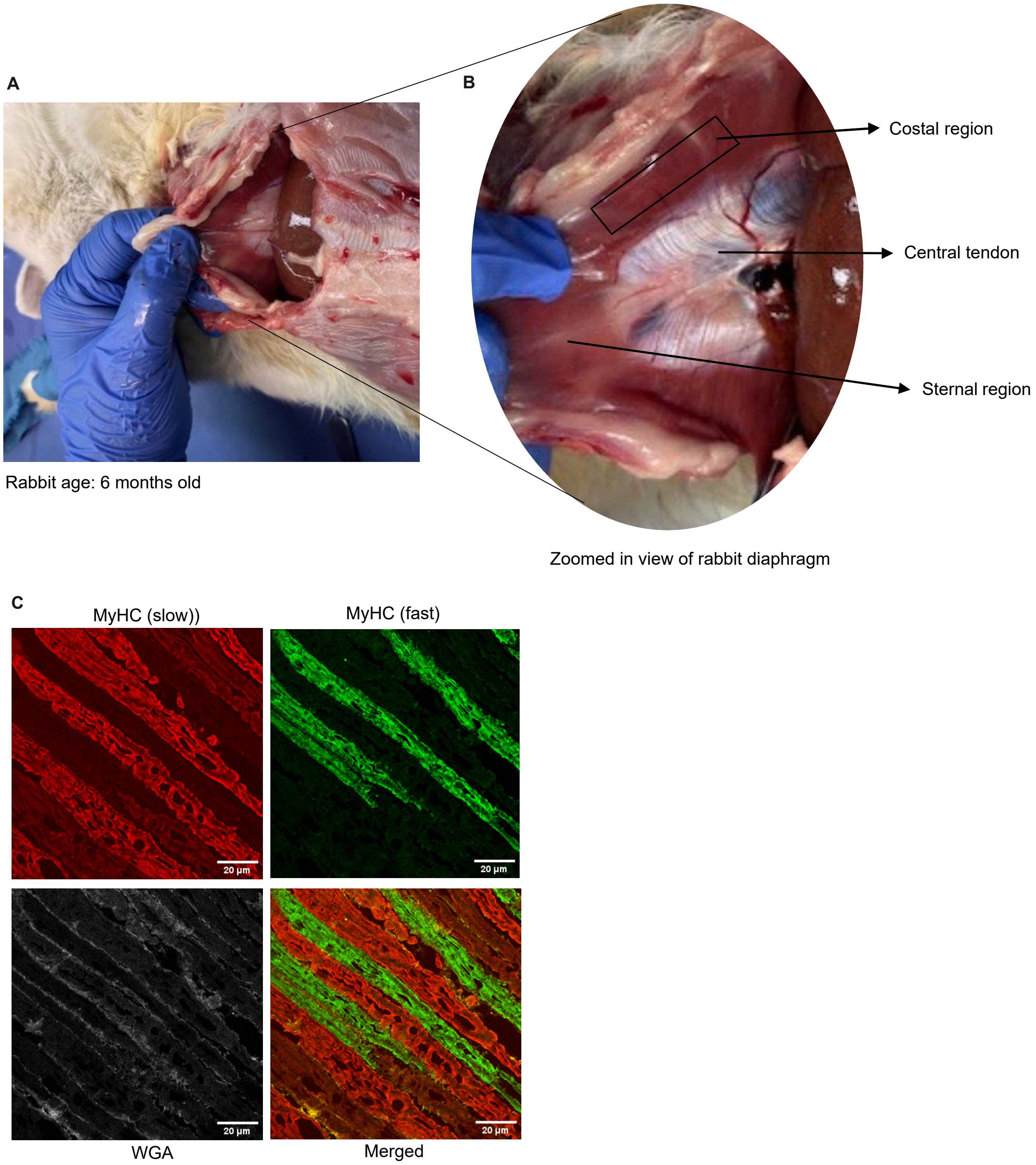


**Fig. S1: Anatomical localization and standardized sampling of the rabbit diaphragm. (A)** Original image of the rabbit diaphragm. **(B)** An enlarged view of the diaphragm identifying the costal and sternal regions and the intervening connective tissue. Tissue collected from the costal region was used for subsequent histological, biochemical, and functional experiments. **(C)** Representative longitudinal cryosections immunolabelled for slow myosin heavy chain (MyHC; red) and fast MyHC (green). Wheat germ agglutinin (WGA; white) delineates individual muscle-fibre boundaries; the merged image shows slow- and fast-MyHC-positive fibres in a diaphragm tissue section.

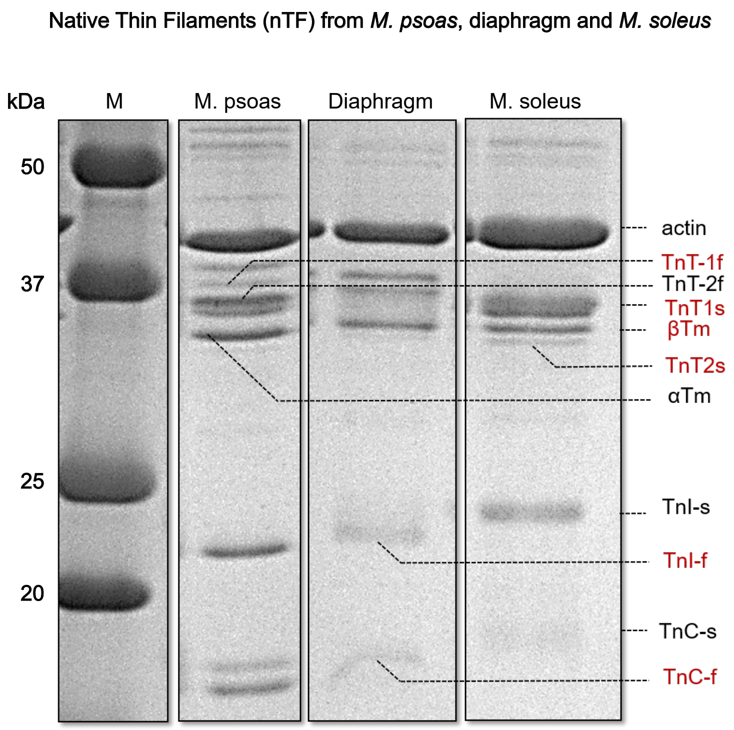


**Fig. S2. Characterization of diaphragm native thin filaments.** Representative SDS-PAGE analysis of native thin filaments isolated from rabbit psoas, diaphragm and soleus muscles. Only the lower part of the gel is shown to highlight the components. All preparations contain actin together with thin-filament regulatory proteins, including troponin T (TnT), troponin I (TnI), troponin C (TnC) and tropomyosin (Tm). nTFs from psoas and soleus muscle tissue preparations display predominantly fast- and slow-skeletal regulatory-protein isoforms, respectively, whereas the diaphragm preparation contains bands corresponding to its mixed slow- and fast-fibre composition (Marked in red). f-fast, s-slow, M, molecular-mass marker.

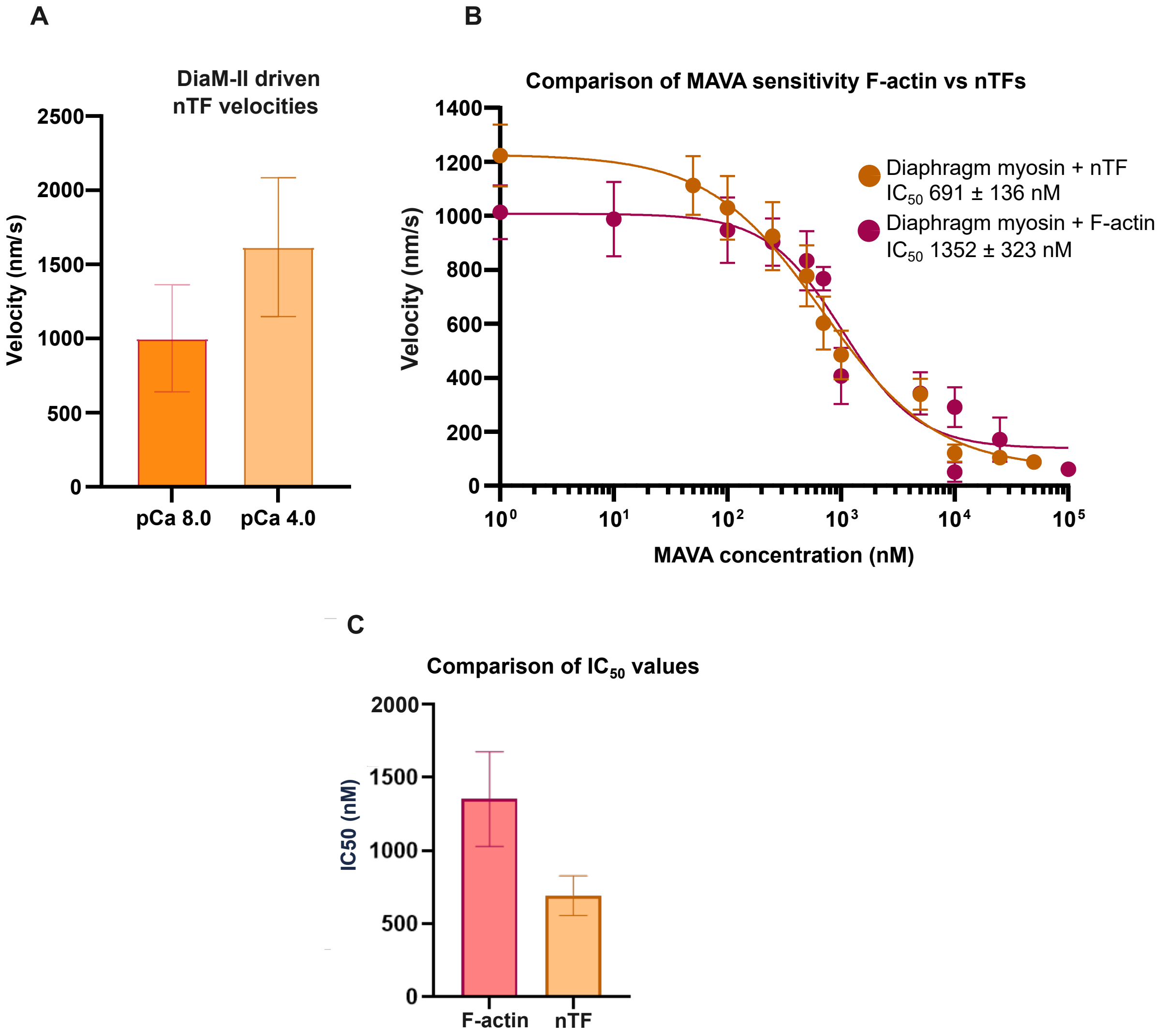


**Fig. S3: Effect of MAVA on diaphragm native thin filaments velocity. (A)** Mean DiaM-II driven gliding velocities of purified native thin filaments at pCa 8.0 and pCa 4.0. **(B)** Comparison of MAVA-dependent inhibition of DiaM-II in the presence of F-actin and native thin-filament. Nonlinear regression fitting - a four-parameter logistic model with variable slope was used to obtain $IC_{50}$ values. **(C)** Comparison of $IC_{50}$ values showing approximately twofold greater MAVA sensitivity with native thin filaments than with purified F-actin. Experiments were performed at 2 mM ATP and 22°C. N = 2-5, n =200-350. Error bars - mean ± SD.

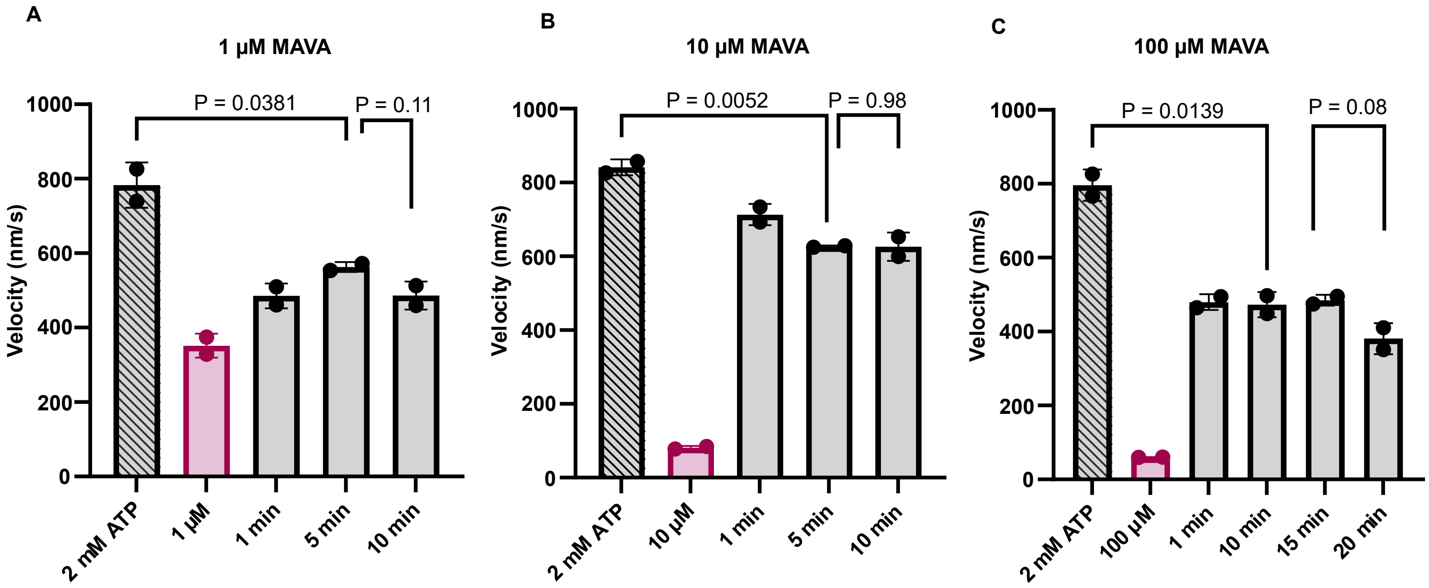


**Fig. S4: Time-dependent recovery of DiaM-II activity following MAVA washout.** Time-dependent recovery following treatment with **(A)** 1 µM MAVA treatment (Red bar) followed by recovery recorded 1, 5 and 10 min after drug washout), **(B)**10 µM MAVA and **(C)**100 µM MAVA. Note that fresh 2 mM ATP containing solution was infused in the chamber before recording the recovery time points. Measurements at 2 mM ATP at 22°C. N =2, n = 100-140. Error bar: mean ± SD. For statistical significance Unpaired two-tailed Student´s t-test was employed for comparison between indicated groups.

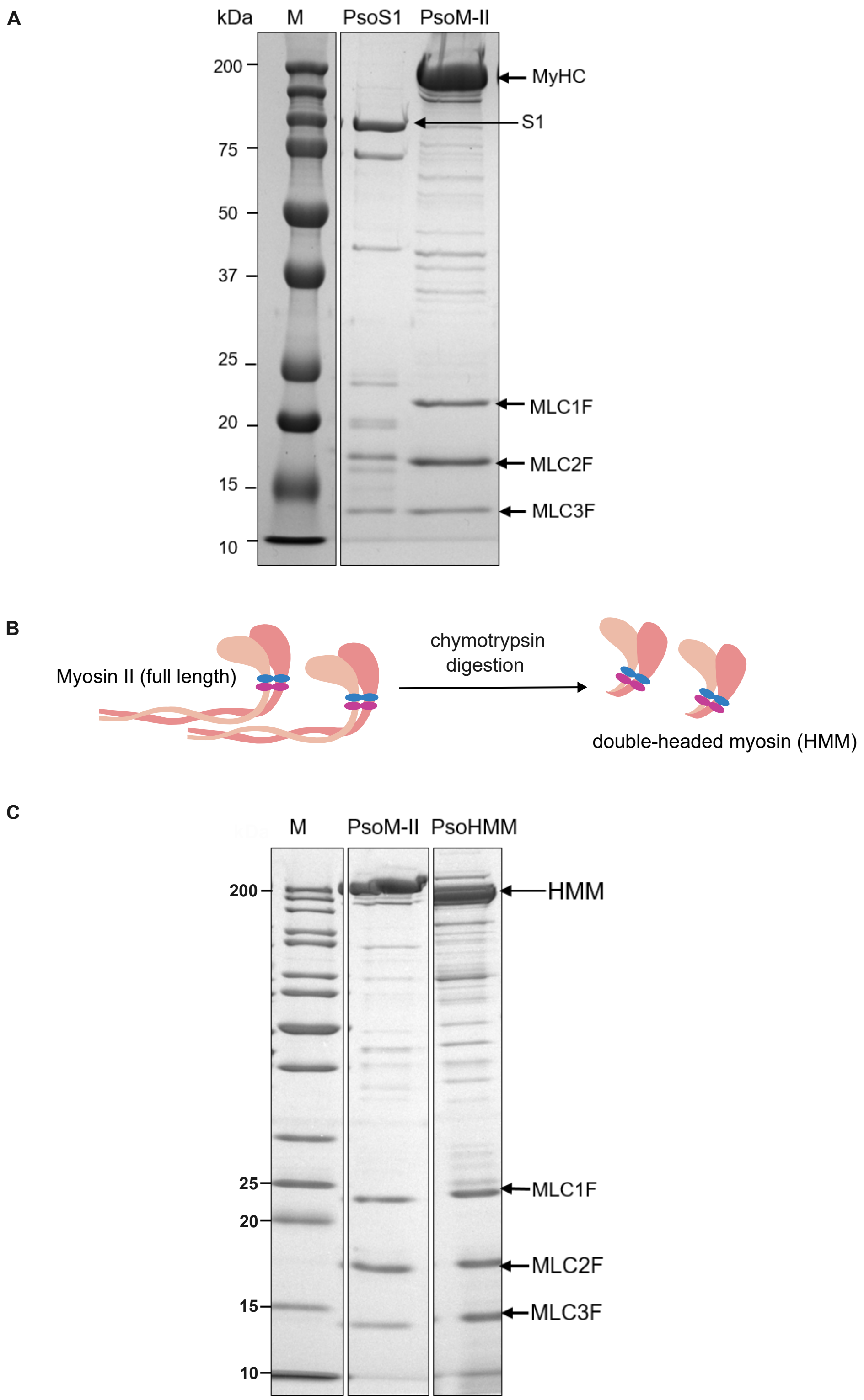


**Fig. S5: PsoS1 and Pso-HMM preparation. (A)** Representative SDS-PAGE analysis of generated PsoS1. The major S1 fragment and light chains are indicated. **(B)** Schematic representation of the generation of psoas heavy meromyosin (Pso-HMM) from full-length rabbit psoas myosin II (PsoM-II) by chymotrypsin digestion. **(C)** Representative SDS-PAGE analysis of Pso-HMM. M, molecular-mass marker. Note that different lanes from the same gels are cropped and assembled together for better visualization.

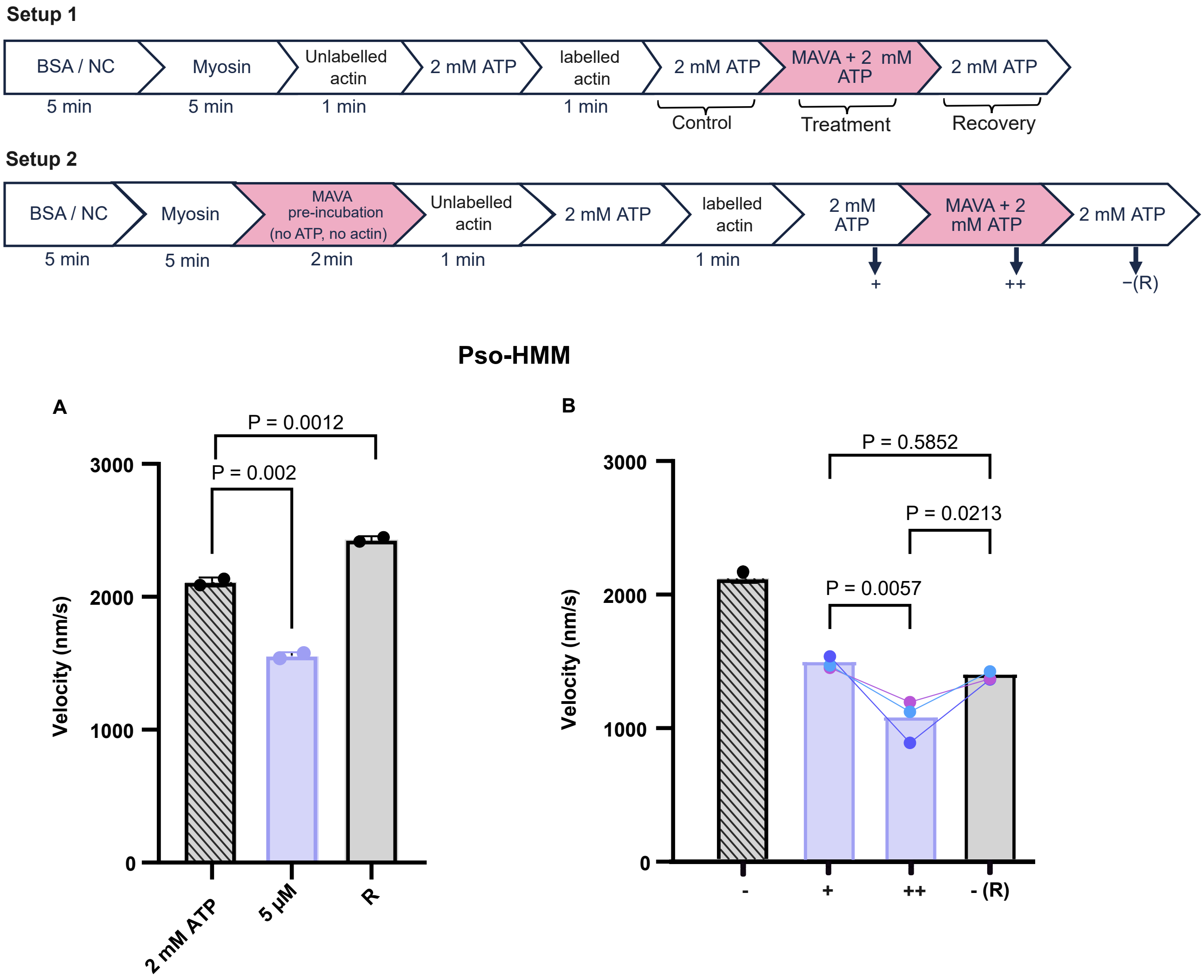


**Fig. S6: Effects of MAVA on double-headed Pso-HMM.** Two experimental setups 1 and 2 are shown as in Figure 7. **(A)** Effect of 5 µM MAVA on rabbit psoas heavy meromyosin (Pso-HMM) using the standard in vitro motility assay following the Setup 1. **(B)** Measurement as per experimental setup 2. The symbols +, ++, and ( -/R) represents the conditions as described in the setups above. - denotes control condition at 2 mM ATP. Connected data points of the same colour represent measurements from the same motility chamber across successive conditions. N = 2-3, n = 100-130. Error bars - mean ± SD. Statistical significance was assessed using an unpaired two -tailed Student´s t-test (setup 1) and ANOVA (setup 2). N = Number of chambers, n = number of actin filaments.

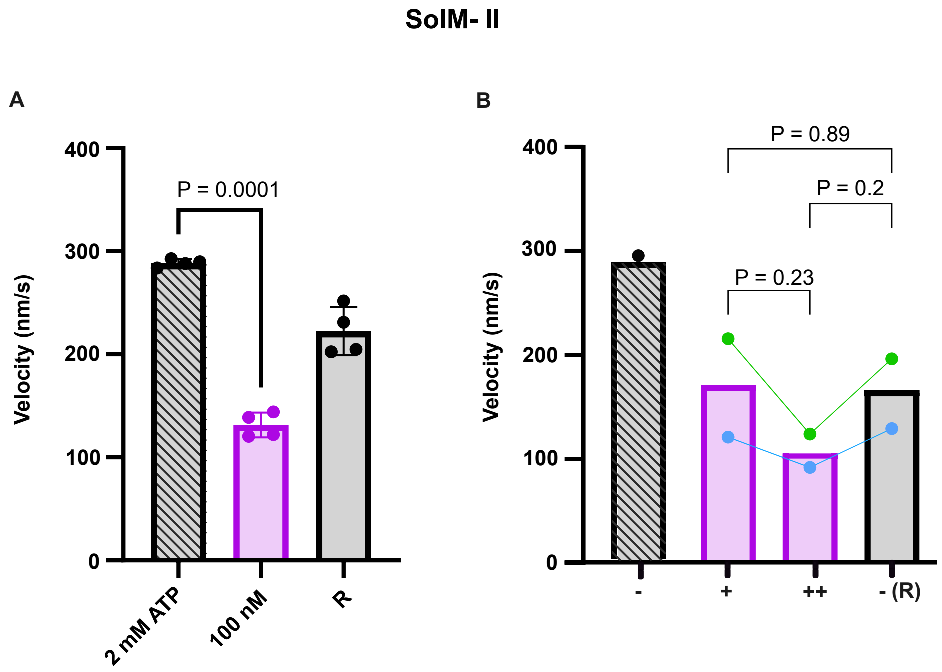


**Fig. S7: Effects of MAVA on SolM-II. (A)** Effect of 100 nM MAVA on SolM-II using the standard in vitro motility assay (setup 1). **(B)** Measurement as per experimental setup 2 as described for Fig. 7 and S6. N = 2-3, n = 100-130. Connected data points of the same colour represent measurements from the same motility chamber across successive conditions. + - after MAVA preincubation, ++ - MAVA in motility buffer, -(R) - recovery. Velocities as as mean ± SD. Statistical significance was assessed using unpaired two-tailed Student´s t-test (setup 1) and ANOVA (setup 2). N = Number of chambers, n = number of filaments.

## βM- II

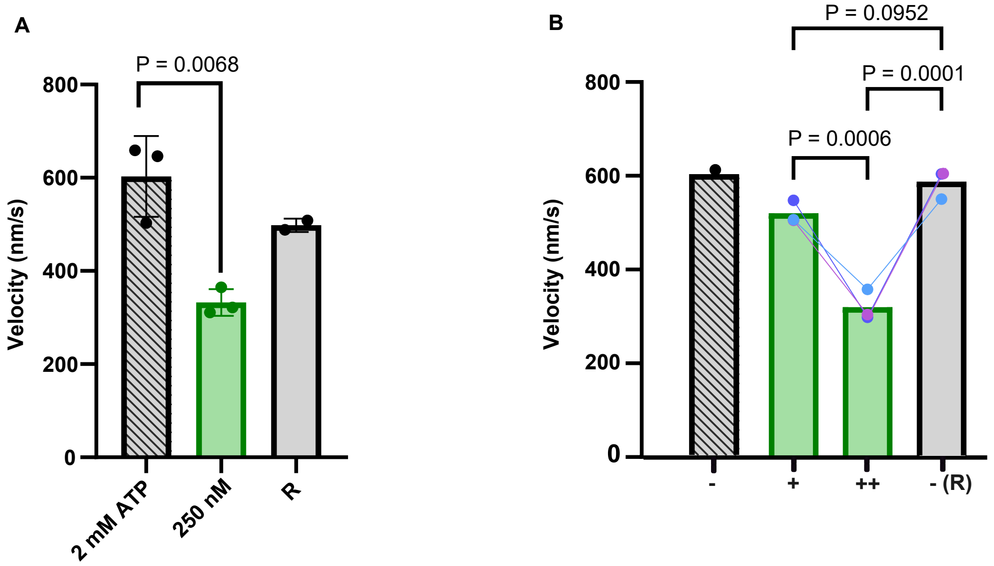


**Fig. S8: Effects of MAVA on βM-II. (A)** Effect of 250 nM MAVA on βM -II using the standard in vitro motility assay (Setup 1). **(B)** Measurement as per experimental setup 2 as described for Fig. 7B and S6. N = 2-3, n = 120-150. Connected data points of the same colour represent measurements from the same motility chamber across successive conditions. + - after MAVA preincubation, ++ - MAVA in motility buffer, -(R) - recovery. Velocities as as mean ± SD. Statistical significance was assessed using an unpaired two-tailed Student´s t-test (setup 1) and ANOVA (setup 2). N = Number of chambers, n = number of actin filaments.

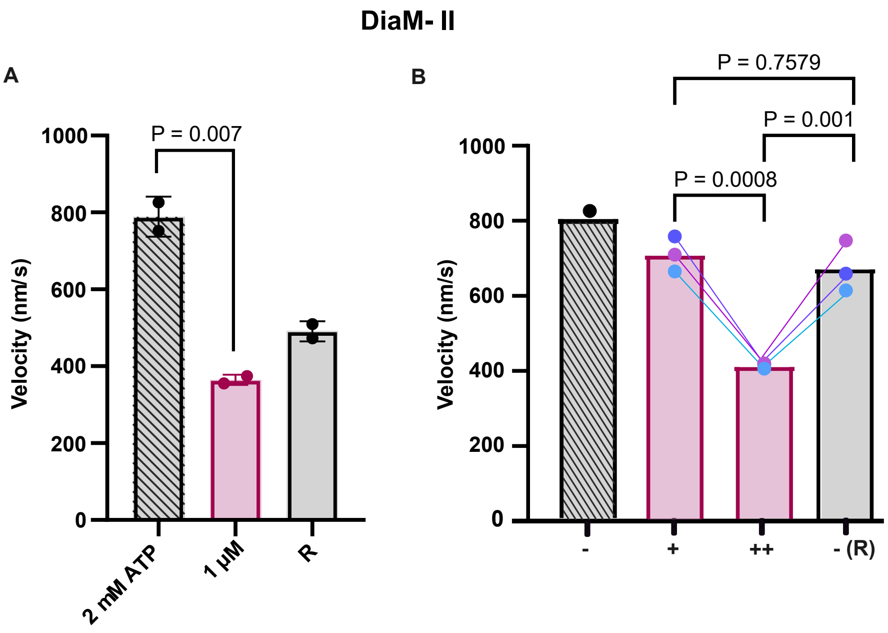


**Fig. S9: Effects of MAVA on DiaM-II. (A)** Effect of 1 µM MAVA on DiaM -II using the standard in vitro motility assay (Setup 1). **(B)** Measurement as per experimental setup 2 as described for Fig. 7B and S6. N = 2-3, n = 100-120. Connected data points of the same colour represent measurements from the same motility chamber across successive conditions. + - after MAVA preincubation, ++ - MAVA in motility buffer, -(R) - recovery. Velocities as as mean ± SD. Error bars - mean ± SD. Statistical significance was assessed using an unpaired two-tailed Student´s t-test (setup 1) and ANOVA (setup 2). N = Number of chambers, n = number of actin filaments.
